# A hybrid boundary element-finite element method to solve the EEG forward problem

**DOI:** 10.1101/2021.12.27.474309

**Authors:** Nasireh Dayarian, Ali Khadem

## Abstract

This article presents a hybrid boundary element-finite element (BE-FE) method to solve the EEG forward problem and take advantages of both the boundary element method (BEM) and finite element method (FEM). Although realistic EEG forward problems with heterogeneous and anisotropic regions can be solved by FEM accurately, the FEM modeling of the brain with dipolar sources may lead to singularity. In contrast, the BEM can solve EEG forward problems with isotropic tissue regions and dipolar sources using a suitable integral formulation. This work utilizes both FEM and BEM strengths attained by dividing the regions into some homogeneous BE regions with sources and some heterogeneous and anisotropic FE regions. Furthermore, the BEM is applied for modeling the brain, including dipole sources and the FEM for other head layers. To validate the proposed method, inhomogeneous isotropic/anisotropic three- and four-layer spherical head models are studied. Moreover, a four-layer realistic head model is investigated. Results for six different dipole eccentricities and two different dipole orientations are computed using the BEM, FEM, and hybrid BE–FE method together with statistical analysis and the related error criteria are compared. The proposed method is a promising new approach for solving realistic EEG forward problems.

## 1 Introduction

Electroencephalography (EEG) is a non-invasive, fast, and inexpensive method to record electrical potentials on the head surface. In neuroscience, it is important to characterize the sources of measured EEG signals and accurately localize them by solving an EEG inverse problem. The EEG inverse problem includes an EEG forward problem using a chosen source model. The solution of the EEG forward problem yields an accurate calculation of the electromagnetic fields. To solve the EEG forward problem, the conductivity profile of the head is modeled, and the relation between the source model and the computed EEG signals is introduced in the EEG lead-field matrix, which can then be used to solve the EEG inverse problem. Then, the source locations and strengths are estimated from the measured EEG signals with the help of the EEG lead-field matrix obtained in the EEG forward model. Consequently, appropriate modeling of the EEG forward problem is an essential prerequisite for the accurate solution of the EEG inverse problem [1, 2]. In other words, the physics of the problem is in the forward model, and errors caused by an inaccurate forward model cannot be corrected while solving the inverse problem.

Two advanced numerical methods called Finite Element Method (FEM) [3], [4] and Boundary Element Method (BEM) [3], [5], [6] are widely used to solve the EEG forward problem. In the FEM, the entire volume is discretized into small elements (tetrahedral elements), and the potentials of all nodes are calculated. The FEM can easily incorporate arbitrary geometries and heterogeneous and anisotropic electrical conductivity of the head tissues [7]-[9]. Unfortunately, the difficulty thorough using FEM is that it causes singularity when using the point dipoles as current dipoles in the EEG forward model, which increases the error of forward solution. However, the current dipoles are widely accepted models for modeling neuronal activities [3], [7], [8].

Some methods have been proposed to improve the behavior of the FEM in singularity cases, e.g. the subtraction method [7], [8], [10], [11] and the direct methods [7], [12]-[14],. The subtraction method has a reasonable mathematical basis for point current dipole models. However, it is computationally relatively expensive and sensitive to conductivity jumps if the source is near them [8], [15]-[17] [10]. On the other hand, The direct FEM approaches such as St. Venant [12] and partial integration [13] are easy to implement and have a much lower computational complexity, so they are very fast [17], [18]. However, the potential distribution strongly depends on the shape of the element at the source location [10], [19]. Among the direct approaches, the St. Venant approach was shown to have the most accurate results for the sources of not very high eccentricity [18]. On the other hand the partial integration approach was shown to have higher stability even at the sources of high eccentricity [19].

On the other hand, the BEM is used for calculating the potentials of surface elements on the interface between compartments generated by a current source in piece-wise homogenous volume. The BEM can construct realistic geometry of piece-wise homogeneous isotropic compartments and solve the EEG forward problem accurately [3]. Also, it has numerical stability and effectiveness compared with differential equation-based techniques [6], [20]. Unfortunately, the BEM formulations can be complicated to model complex geometry such as inhomogeneity, anisotropicity and surfaces with holes [3], [6], [21]. Also, the BEM produces dense matrices that cause high computational cost compared with FEM [21].

In order to benefit from the advantages of both BEM and FEM, some coupled boundary element (BE)–finite element (FE) methods have been proposed in electromagnetic and biomedical problems [22]-[26]. In [22], a new high-order cubic Hermite coupled FE/BE method has been proposed only for an isotropic three-layer spherical and realistic head model, and generalized Laplace’s equation had been solved. Also, a hybrid BE–FE method has been applied to the 2D forward problem of electrical impedance tomography (EIT) [23] and has been used for modeling Diffusion equations in 3D multi-modality optical imaging [24]. A 3D coupling formulation was presented in [25] for solving the EEG forward problem iteratively. A domain decomposition (DD) framework was used to split the global system into several subsystems with smaller computational domains. Then, for each subsystem, one of the methods (BEM or FEM) was used. Several iterations were needed to solve the forward problem on the global system, and a relaxation parameter at each interface was compulsory to ensure convergence. The relaxation parameters were set manually, and an inappropriate value of these parameters would make the scheme diverge. Furthermore, three-layer concentric sphere models considering both the isotropic and anisotropic conductivity of the skull layer with dipoles of six locations and three orientations were modeled. No realistic head model was investigated. The coupling process was very time-consuming because the BEM and FEM ran iteratively until the relative residuals reached below a properly set value (6×10^-5^).

In [26], a hybrid BE–FE method, which directly combines the two BE and FE methods, has been proposed to solve the forward problem of EIT for a 3D cylindrical model of the human thorax. It should be noted that the EEG forward problem is completely different from the EIT forward problem regarding equations and boundary conditions. Thus, we must reformulate equations and extend them to be suitable for applying to the EEG forward problem. The advantage of using such a hybrid BE–FE method for solving the EEG forward problem is that the isotropic and homogeneous subregions containing the current dipoles can be modeled by the BEM and the other subregions (the inhomogeneous or anisotropic subregions or those without current dipoles) can be modeled by the FEM. Also, this method solves the global system in one step without any iteration. Consequently, it is expected that the application of the hybrid BE–FE method increases the accuracy of the EEG forward solution and consequently helps to more accurate localization of brain sources.

In this paper, BEM, partial integration FEM (PI-FEM), and hybrid BE–FE method, are employed to solve the EEG forward problem. To validate the hybrid BE–FE method in solving the EEG forward problem of isotropic multi-compartment media, we will use an isotropic piece-wise homogeneous three-layer spherical head model (brain, skull and scalp), which has an analytical forward solution, and the results will be compared with those of BEM and PI-FEM. To validate the hybrid BE–FE method in solving the EEG forward problem of anisotropic multi-compartment media, we will use an anisotropic three-layer spherical head model (brain, skull and scalp) in which the conductivity of the skull will be considered anisotropic and compare the results with those of PI-FEM. Since the cerebrospinal fluid (CSF) layer highly affects the scalp potentials [27], [28], we will also investigate the performance of the hybrid BE–FE method compared with PI-FEM when considering the fourth layer for CSF and the anisotropic conductivity of the skull.

In [29], it is considered that the human skull has a three-layered sandwich structure, consisting of a hard (compact) bone enclosing another layer of soft (spongy) bone which is not equally thick everywhere. In cases that accurate identification these two tissue types (e.g., from MRI) and exact knowledge of their conductivity values is an issue, an approximated model of the skull as a single-layer of homogeneous and isotropic/anisotropic profile with optimized value of conductivity/anisotropy ratio can be used [30]. In this paper, the technique introduced in [30] is used. It is noteworthy that our proposed method is able to model a complex profile of skull.

Because the conductivity uncertainties of head layers (especially skull and brain) have a significant influence on the EEG forward solution [31], we will repeat the simulation of each spherical head model 50 times with different realizations of conductivity of layers followed by statistical analysis to demonstrate the effect of conductivity uncertainties on the EEG forward solution.

Finally, the hybrid BE–FE method and PI-FEM will be compared on a four-layer realistic head model in which the conductivity of the skull will be anisotropic.

This paper is organized as follows: In Section 2, the mathematical model for the EEG forward problem and its numerical solutions using BEM, PI-FEM, and hybrid BE–FE method will be formulated. In Section 3, the performance criteria for validation will be described. In Section 4, the results of simulated spherical and realistic head models will be reported. In Section 5, the results will be discussed. Finally, in Section 6, the paper will be concluded, and some future works will be proposed.

## 2 Mathematical model

In this section, first, the mathematical model of the EEG forward problem is introduced, and then the BEM, PI-FEM and hybrid BE–FE method are formulated to solve the EEG forward problem. Finally, we explain how to generate mesh for each method.

### 2-1 EEG forward problem

The EEG forward problem entails calculating the electric potential *φ* on the scalp surface S (see Figure 1). These potentials are generated by current dipoles within the head volume R. Therefore, since the relevant frequencies of the EEG spectrum are below 100 Hz, the quasi-static approximation of Maxwell’s equations is used to estimate the electric potentials over the scalp with homogeneous Neumann boundary conditions as follows [32]:

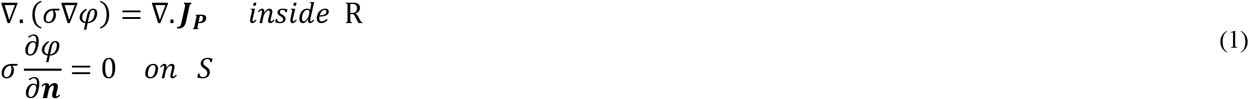

where *σ*: ℝ^3^ → ℝ^3^ denotes conductivity tensor of tissue conductivity in R and ***J_P_*** denotes the primary current density of the brain source. Also, ***n*** is the outward unit normal vector at the surface *S* [3], [32]. In this manuscript, vector quantities are denoted by bold characters.

**Figure 1:**
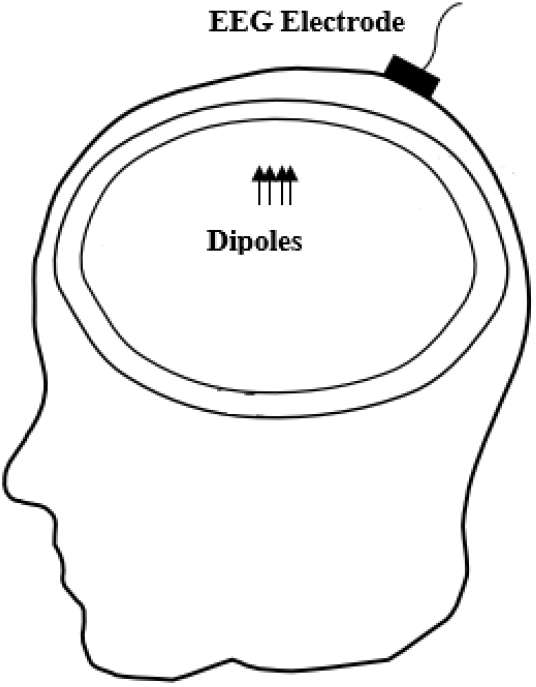
EEG forward model. Electric potentials recorded by EEG electrodes are generated by dipoles within the head volume. A sample EEG electrode placed on the scalp is only shown.

The primary current density ***J_P_*** is commonly modeled as two delta functions at the current source position ***r***_2_(*x*_2_, *y*_2_, *z*_2_) and the current sink position *r*_1_(*x*_1_, *y*_1_ *z*_1_) with the current source density *I* as follows [14]:

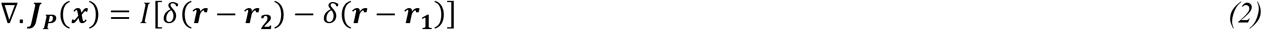

### 2-2 Boundary element method (BEM)

In the BEM, the reciprocal relation is applied to derive a boundary integral equation for the boundary value problem (1) as given by [33]

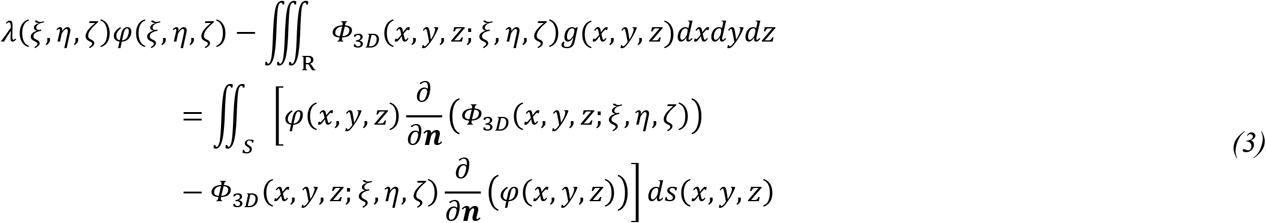

where 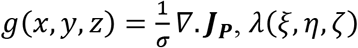 is a characteristic function of the domain *R, σ* is the conductivity of domain that must be constant and isotropic. The function *Φ_3D_*(*x,y,z;,ξ,η,ζ*) is the fundamental solution of the three dimensional Laplace’s equation and is given by [33]

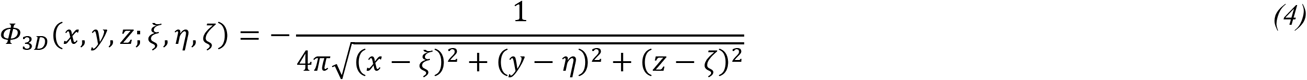

Consider the surface of a region to be discretized to *N*triangular elements. The potential *φ*(*x,y, z*) and its normal derivative 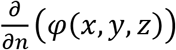 are approximated as constant values over each element as follows:

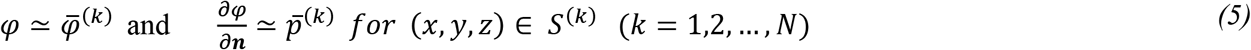

where 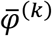 and 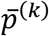 are the average values of *φ* and 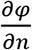 on the centroid point of the k^th^ surface element, *S*^(*k*)^ Using these approximations, (3) is simplified as

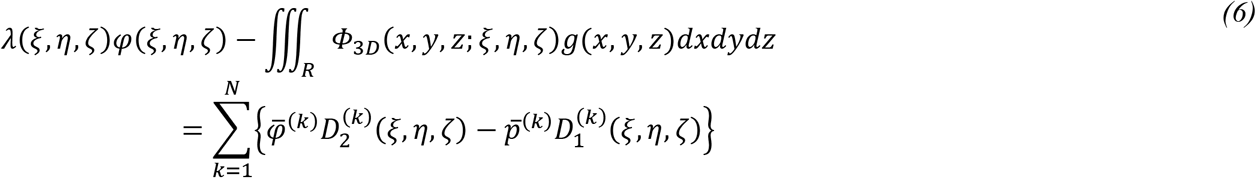

where

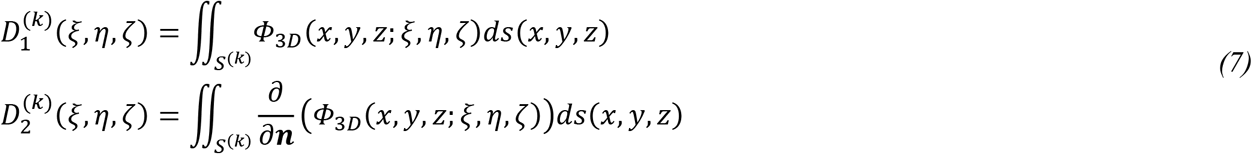

Let (*ξ,η,ζ*) in (6) be given consecutively by the centroid point of *S*^(1)^, *S*^(2)^,..., *S*^(*N*)^. Consequently, (6) can be rewritten as [33]

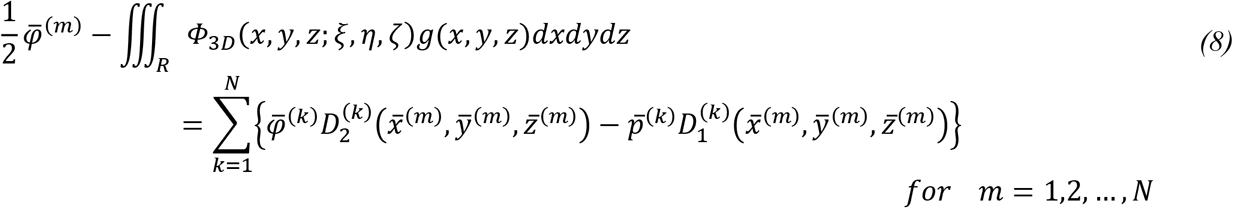

where 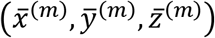 is the centroid point of the element ***S***^(*m*)^. On the typical element *S*^(*k*)^, either 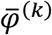 or 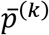 is known. Thus (8) constitutes a system of *N* linear algebraic equations including *N* unknowns for a one-layer homogenous region as given by [33]

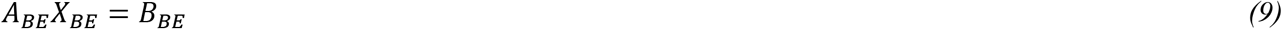

where *X_BE_* is the column vector (column matrix) including both 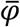 and 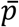 for each element, *A_BE_* is the coefficient matrix and *B_BE_* is the column vector containing values of boundary condition and the current dipoles information. Equation (9) describes the matrix form of the BEM for just one homogenous subregion.

In order to implement BEM for multi-layer piece-wise homogeneous media, we used the same approach as that of [26]. Hereby, we describe that method for a three-layer spherical head model with nested regions of constant conductivity (see Figure 2). First, all interfaces are discretized to *N* triangles, and then the first subregion (brain) is represented by (1). Other subregions (skull and scalp) are represented by the Laplace equation. The first and second boundaries of each subregion are assumed to be Dirichlet and Neumann conditions, respectively. Considering the air/scalp interface, the boundary condition of the second boundary of the scalp is of Neumann type. The system of linear algebraic equations for each subregion is obtained as follows:

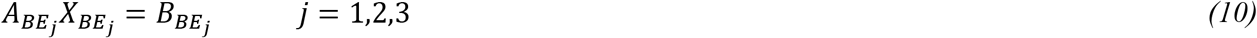

where *X_BE_j__* is the column vector including both 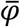 and 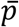 for each element at the *j*^th^ subregion. Assembling the equations corresponding to all subregions, the same algebraic equation as (9) is derived. Then, continuity conditions at the interface between two adjacent subregions should be applied to the resulting equation. The continuity condition for the electric potential and continuity condition for the normal component of the current density at the interface surface *S_BE-BE_* are given as follows:

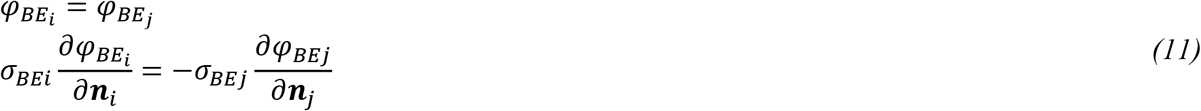

where (*φ_BE_i__*, *σ_BE_i__*) and (*φ_BE_j__, σ_BE_j__*) are the (electric potential, conductivity) of each element on the interface between two adjacent subregions Ω*_BE_i__* and Ω*_BE_j__*, respectively. After the continuity condition for the electric potential and the normal component of the current density, the electric potential and its normal derivation of surface elements are computed by using Gaussian elimination method.

**Figure 2:**
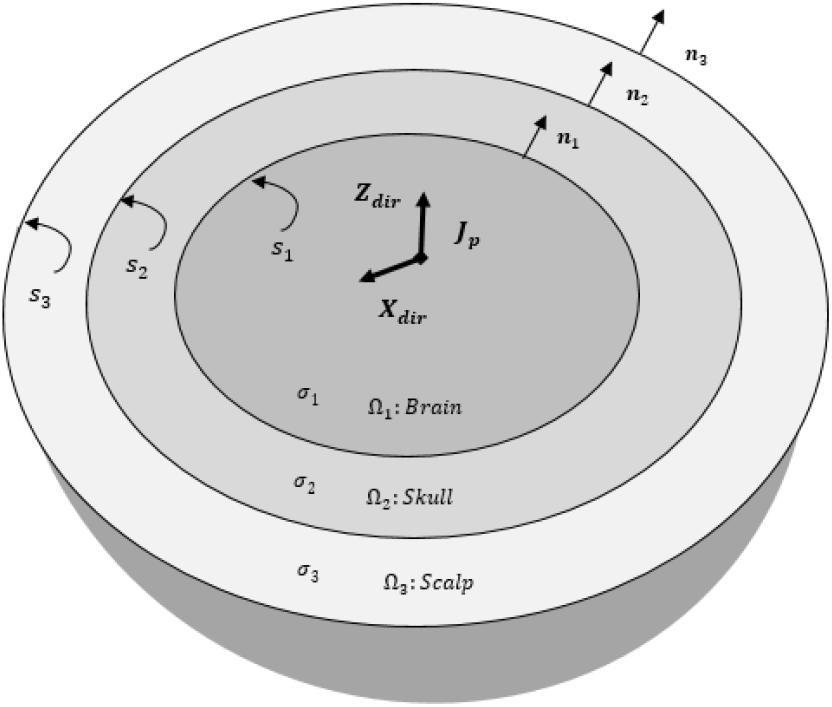
The piece-wise homogenous three-layer spherical head model (brain, skull, and scalp).

### 2-3 Finite element method using Partial Integration approach (PI FEM)

In the FEM, the functional *F*(*φ*) derived from the Rayleigh-Ritz method, a variational method, is minimized to solve the boundary value problems. Equation (1) can be written in Cartesian coordinates with homogeneous Neumann boundary conditions as follows [34].

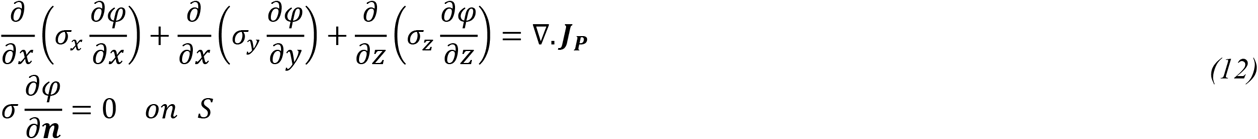

Consequently, the functional is defined as [34]:

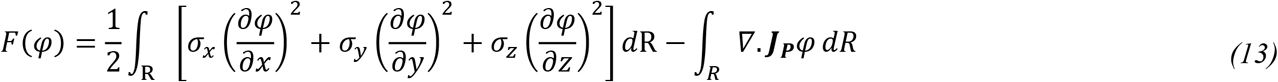

where *σ_x_, σ_y_* and *σ_z_* are respectively the conductivity along x, y and z axes that are constant and equal to each other in isotropic media but unequal to each other in anisotropic media.

The first step of the FEM is the discretization of the regions into a number of tetrahedral elements. The unknown potential *φ^e^* at any point within each tetrahedral element can be approximated as:

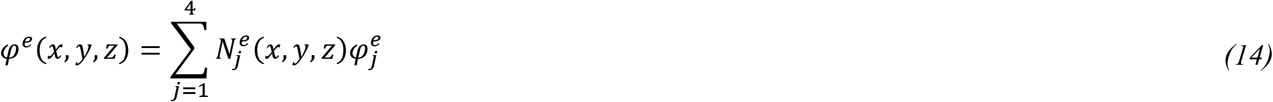

where the interpolation functions 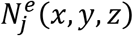 are given by

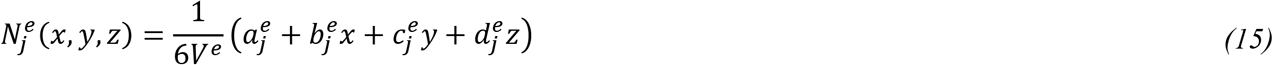

where 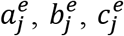 and 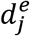 are constants and are determined from the coordinates of the nodes of elements. By minimizing the functional *F*(*φ*) in (13) for each element and assembling all elements in the whole volume, and using the partial integration(PI) approach to model the current dipoles [13], [35], the final set of equations can be written as follows [34].

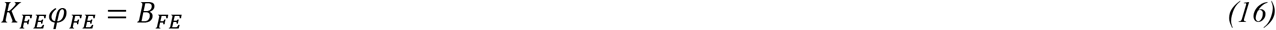

where *K_FE_* is the coefficient (stiffness) matrix, which is a function of nodal coordinates and conductivity of each element. *φ_FE_* is the column vector of the unknown electric potential of the nodes and *B_FE_* is the source column vector contributed by the dipoles that has non-zero entries for the set of nodes of the elements that contain the dipoles. After the Neumann boundary condition, given in (1) is applied to (16), the electric potential is computed by using quasi-minimal residual method.

### 2-4 Hybrid BE–FE method

The hybrid BE–FE formulation consists of both FE and BE formulations. It is implemented by combining (9) with (16) and applying boundary conditions in each interface. First, the BEM is used to represent the Poisson equation (1) at the brain subregion (BE region) considering the Dirichlet boundary condition. Then FEM is used to represent the Laplace equation (∇. (*σ*∇*φ*) = 0) at other subregions (FE regions) considering Neumann condition at skull/brain and air/scalp interfaces for the three-layer spherical head model, and CSF/brain and air/scalp interfaces for the four-layer spherical head model to derive (16). The values of boundary conditions on each interface are unknown. Assembling (9) and (16) gives

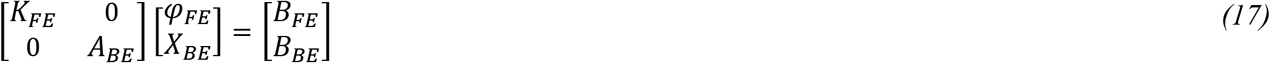

To solve (17), the value of boundary condition at the BE–FE interface must be applied but it is unknown. To apply boundary condition on a BE–FE interface *S_BE-FE_*, the potential on *S_BE-FE_* computed from both BE region and FE region must be the same. Since the potential on a surface BE element is constant, but in a FE element is linear, in order to equalize those potentials on the BE–FE interface, one may take the average of three FE nodal potentials and obtain the following approximation for the BE surface element potential as [26]

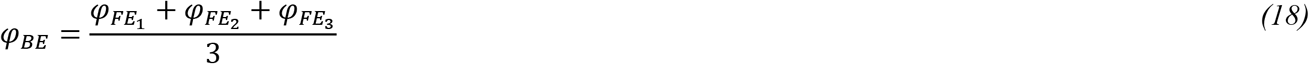

where *φ*_*FE*_1__, *φ*_*FE*_2__, and *φ*_*FE*_3__ are FE nodal potentials at each element in *S_BE-FE_*. The continuity condition for the normal component of the current density at *S_BE-FE_* yields (19) [26]

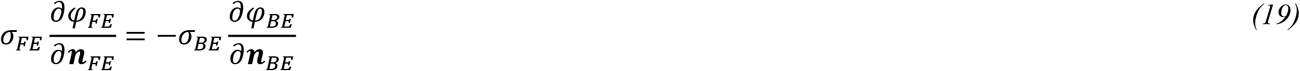

Applying (18) and (19) to (17), the matrices *K_FE_, A_BE_, B_FE_*, and *B_BE_*, are modified as 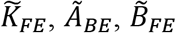, and 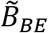, respectively; and the resulting equation is obtained as

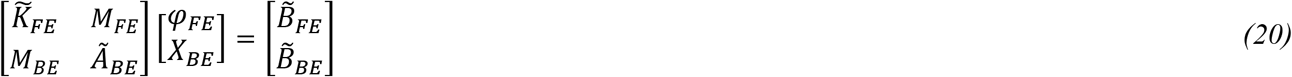

where *M_BE_* and *M_FE_* are sparse matrices constructed as a result of applying (18) and (19), respectively. Using Gaussian elimination method, (20) is solved for *φ_FE_* and *X_BE_*.

To use the hybrid BE–FE method, two important points should be considered:

First, when an anisotropic piece-wise homogenous multi-layer medium is modeled, the layer(s) containing dipoles must be modeled with the BEM, and the layers with anisotropic conductivity must be modeled with the FEM. Other layers can be modeled with each of FEM or BEM depending on our goals, such as less time and memory consumption, and computational complexity. So in this paper, we used the FEM for other layers. Also, we will show the performance of the hybrid BE–FE method when only one layer, including current dipoles is modeled with the BEM, and we compare this performance with the performance of FEM.

Second, we used linear elements in the FEM. Thus, to couple BEM and FEM elements, we need constant triangular elements or continuous linear triangular elements in BE domain to use (18) and (19). We prefer to use constant elements in the BEM because solving forward problem with constant elements is more accurate than continuous linear elements [33].

It should be noted though the discontinuous linear element is more accurate than constant and continuous linear elements, it cannot be used to couple with the FEM in our method and because in our method, we need the constant potential in the surface BE elements or the vertex’s potentials of BE elements to use in (18).

### 2-5 Tetrahedral mesh generation

To implement the FEM and BEM, the entire region is first discretized into tetrahedral volume elements for the FEM, and then the required surface triangular meshes in the BEM are prepared from the data of tetrahedral elements of the entire region. For mesh extraction, we use ISO2MESH [36] that provides us with accurate mesh volume and surface elements.

In the hybrid BE–FE method, the BE regions and the FE regions are discretized using a triangular surface mesh and a tetrahedral volume mesh, respectively. In order that the BE and FE regions in the hybrid method have the same boundary surface elements, the entire volume domain is first discretized by irregular tetrahedral volume elements, and then irregular triangular surface elements for the BE regions are extracted from the data of the tetrahedral elements. It is noteworthy that the FEM uses the entire tetrahedral volume elements.

In the realistic head model, we can leverage on FieldTrip software to obtain an automatic segmentation of the brain, CSF, skull, and scalp [37]. By using FieldTrip, the surface boundaries of these four structures are extracted. Then, based on the boundaries, a finite element model can be constructed by ISO2MESH [36].

## 3 Validation method

In this section, first, the validation platform is described. Then error criteria used to validate our approach are introduced.

### 3-1 Validation platform

To validate the proposed hybrid BE–FE method and compare it with the FEM and BEM, we will first simulate an isotropic, piece-wise homogeneous three-layer spherical head model with radial and tangential dipoles of six different eccentricities. Then, we will repeat the same simulation but with an anisotropic layer (skull) to show the performance of FEM and the proposed hybrid BE–FE method when the skull is modeled as a layer of homogenous and anisotropic profile. Afterward, we will add a fourth layer for CSF to the previous anisotropic model. This layer has an important role in distributing the currents in the head model and scalp potentials [27], [28]. Next, we will repeat the same procedure to assess the performance of the hybrid BE–FE method on this anisotropic and piece-wise homogeneous four-layer medium. For each of the above spherical head models, we will simulate 50 realizations using randomly chosen conductivity values from realistic intervals to assess the precision of the hybrid BE–FE method. This allows us to gain a statistical overview of the precision of solving EEG forward problem with regard to the electromagnetic properties of layers. Finally, we will compare the proposed hybrid BE–FE method with the FEM on an anisotropic piece-wise homogeneous four-layer realistic head model.

In each of the above models, unit radial (along z-axis) and tangential (along x-axis) dipoles of six different eccentricities will be considered. For each dipole, the source eccentricity is defined as the percentage of the Euclidian distance between the dipole location and the sphere center divided by the radius of the innermost shell [38]. In our implementations, the most eccentric dipole has an eccentricity of 98%.

In this paper, for spherical head models, we use the analytical solution given in [39] that calculates the electric potential on the outmost surface (scalp) of isotropic/anisotropic multi-layer spherical head model generated by a dipole inside the innermost layer (brain). For the realistic head model, we use a refined model of the FEM to obtain a reference FE solution because an analytical solution is unavailable for real models.

It is noteworthy that in all of our simulations, in order to have fair comparisons between the accuracy of the FEM, BEM and hybrid BE–FE method, we choose the mesh resolution that their computation times are nearly the same.

### 3-2 Error criteria

The error between the numerical solution and analytical solution can be obtained by Relative Difference Measure (RDM) and the Magnitude ratio (MAG) [3], [40]. These measures are respectively defined as follows:

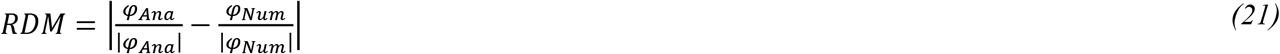

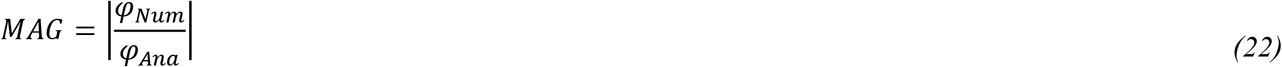

where *φ_Ana_* and *φ_Num_* are analytical and numerical solutions, respectively and | | denotes the square root of Euclidean distance. In this paper, *φ_Ana_* is calculated by using the analytical solution given in [39] for spherical head models and the reference FE solution obtained from the refined model of the FEM for the realistic head model.

## 4 Simulation Results

In this paper, the performance of the hybrid BE–FE method to solve the EEG forward problem will be assessed using both spherical and realistic head models. In the spherical models, the solution will be evaluated on all outer boundary (scalp) nodes instead of the small number of them so that the results are nearly independent of the choice of electrode positions [4]. On the other hand, for the realistic head model, the results will be assessed on 90 positions of the scalp surface.

### 4-4 Example I: Isotropic piece-wise homogenous three-layer spherical head model

To compare the performance of hybrid BE–FE method with the BEM and PI-FEM, a numerical validation will be performed using a three-compartment (brain, skull and scalp) spherical head model as shown in Figure 2 with parameters indicated in Table 1 [31], [41]. The optimized anisotropy ratio in Table 1 is defined as the ratio of radial conductivity (*σ_r_*) to tangential conductivity (*σ_t_*) of the skull [30]. In this simulation, the PI-FEM mesh has 6808 nodes and 34428 tetrahedral volume elements, the BEM mesh has 3092 triangular surface elements, and the hybrid BE–FE mesh has 2578 nodes, 10688 tetrahedral volume elements and 1894 triangular surface elements.

**Table 1:** Parameters of the concentric three-layer spherical head model [31], [41]. The conductivity values, radius and optimized anisotropy ratio are based on [31], [41] and [30], respectively.

| Tissue | Brain | Skull | Scalp |
| --- | --- | --- | --- |
| Outer shell radius (cm) | 8.0 | 8.6 | 9.2 |
| Conductivity interval (S/m) | 0.2200-0.6700 | 0.0016-0.0330 | 0.2800-0.8700 |
| Optimized anisotropy ratio ($\sigma_r/\sigma_t$) | - | 0.0093: 0.015 | - |

Figure 3 and Figure 4 show boxplots of RDM and MAG of PI-FEM, BEM, and hybrid BE–FE method for isotropic and piece-wise homogeneous three-layer spherical head models versus different eccentricities of the dipole for radial and tangential dipoles, respectively. For each model, 50 realizations were simulated by randomly chosen conductivities from intervals shown in Table 1. Also, the mean and standard deviation of RDM and MAG and P-values of Wilcoxon signed-rank tests are reported in Table 2. It should be noted that some datasets didn’t pass the Gaussian test (P-value<0.05). For this reason, we used Wilcoxon signed-rank test to calculate P-values. The significant differences are shown as gray in Table 2.

**Figure 3:**
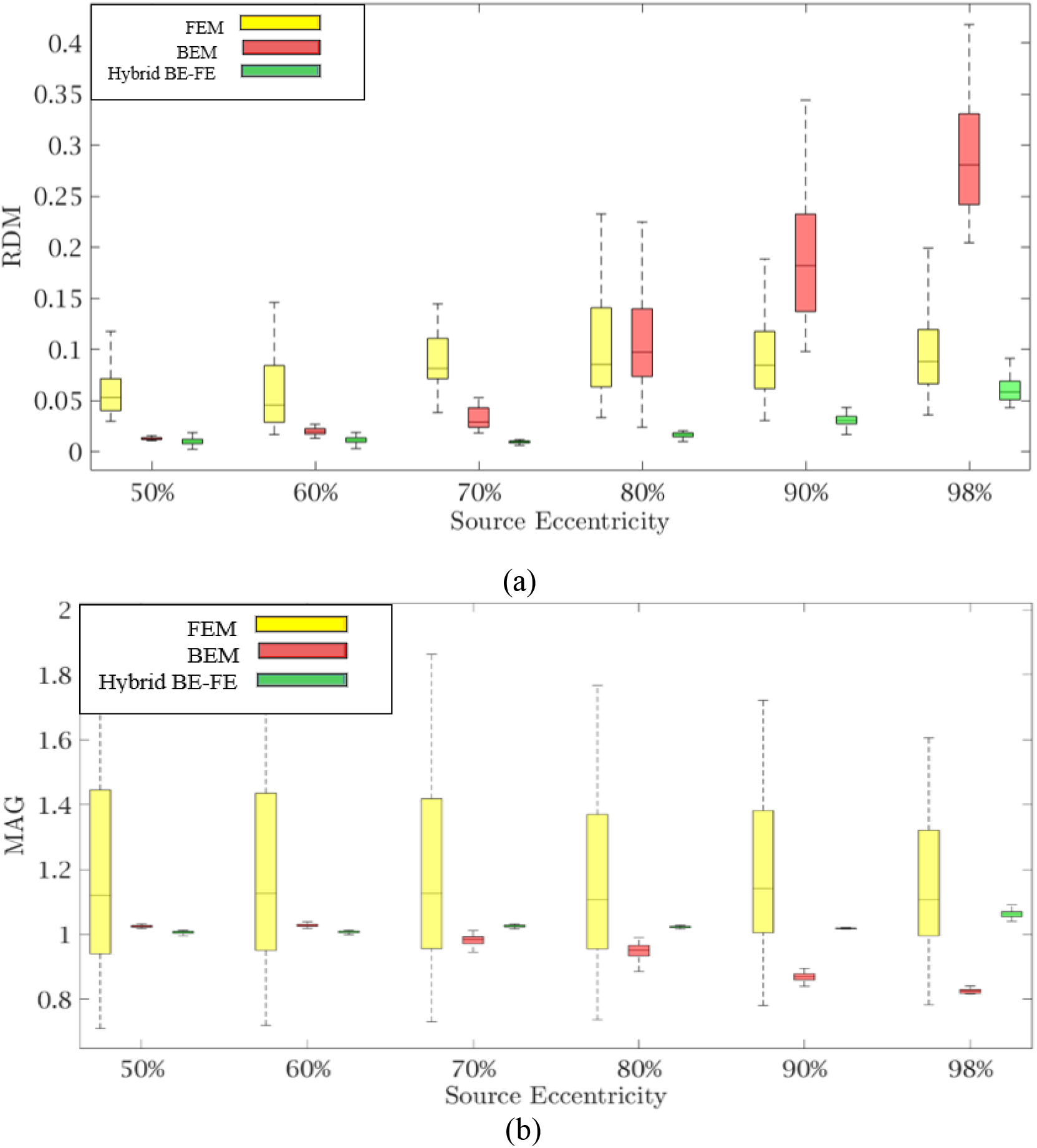
Example I: Isotropic piece-wise homogenous three-layer spherical head model for radial dipole orientation (z-axis), (a) RDM and (b) MAG boxplots of PI-FEM, BEM, and hybrid BE–FE method at six different source eccentricities.

**Figure 4:**
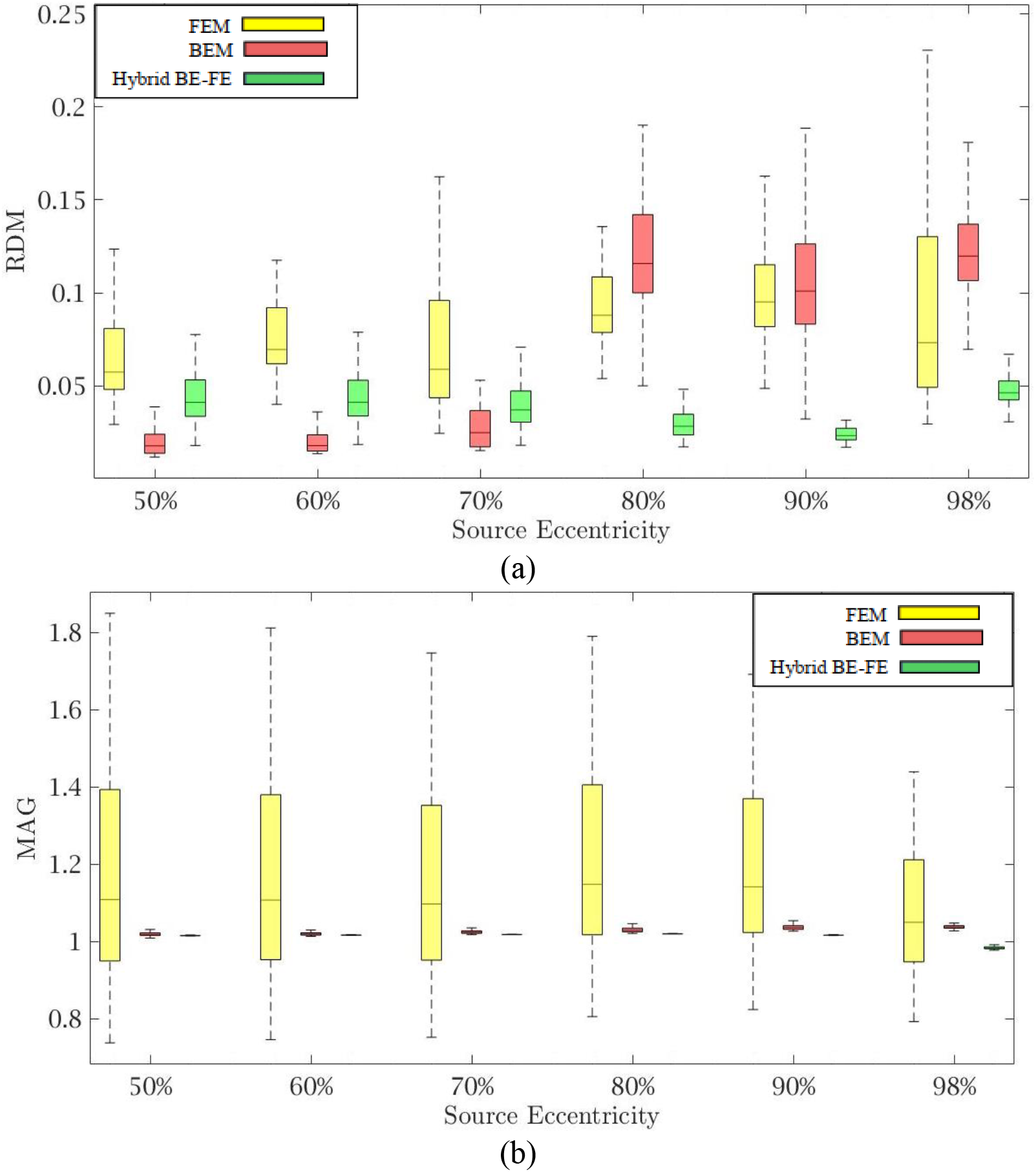
Example I: Isotropic piece-wise homogenous three-layer spherical head model for tangential dipole orientations (x-axis), (a) RDM and (b) MAG boxplots of PI-FEM, BEM, and hybrid BE–FE method at six different source eccentricities.

**Table 2:**
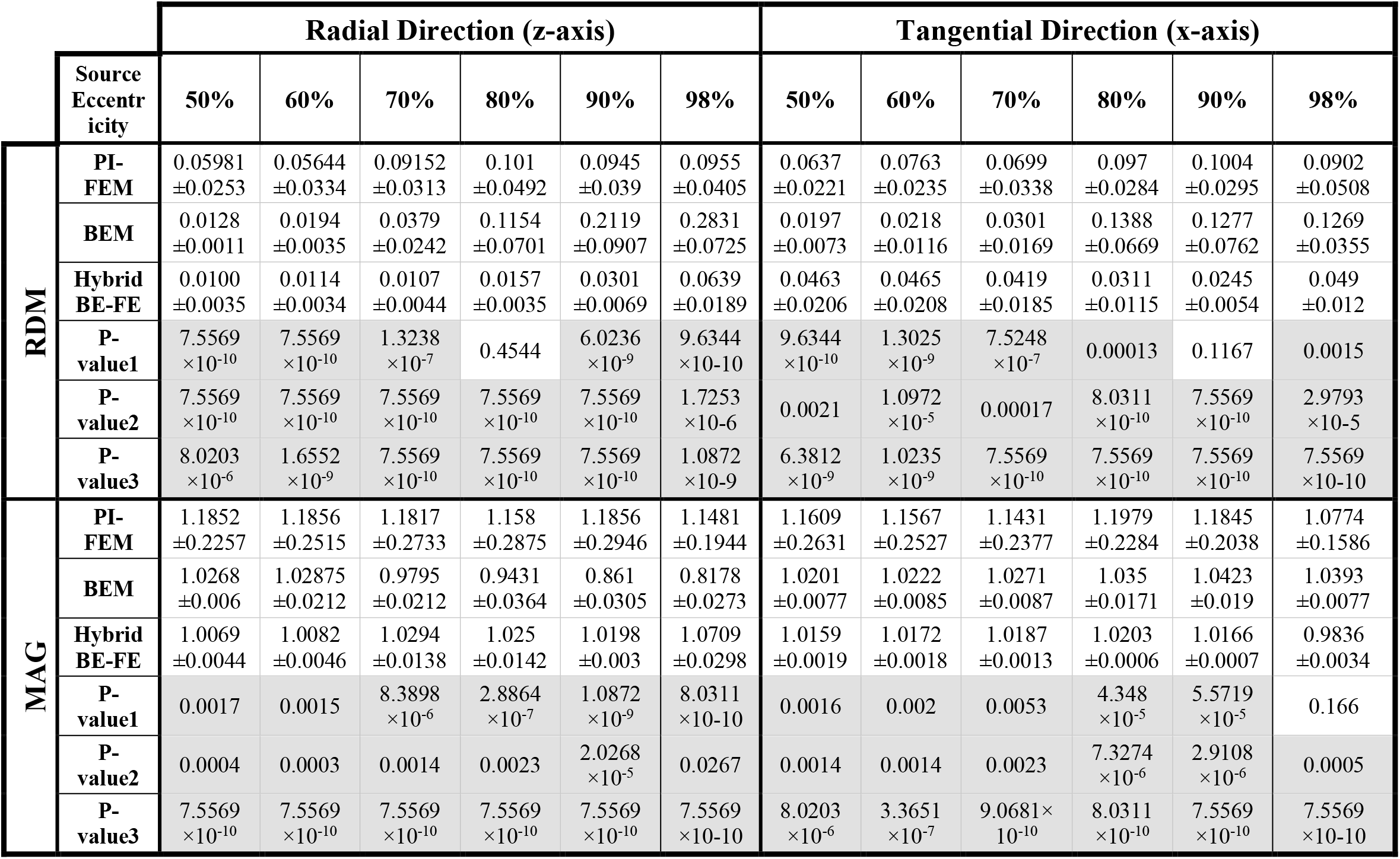
Example I: Isotropic piece-wise homogeneous three-layer spherical head model, mean ± std and P-value of Wilcoxon signed-rank test of 50 realizations of RDM and MAG obtained from PI-FEM, BEM and hybrid BE–FE method for dipoles of six different source eccentricities and both radial and tangential directions. P-value1 corresponds to comparing the results of PI-FEM, and BEM, P-value2 corresponds to comparing the results of PI-FEM and hybrid BE–FE method, and P-value3 corresponds to comparing the results of BEM and hybrid BE–FE method. The significant results are shown as gray.

Comparing the PI-FEM, BEM, and hybrid BE–FE method with regard to the RDM for isotropic piece-wise homogeneous three-layer spherical head model (Figure 2) shows that the hybrid BE–FE method significantly outperforms the PI-FEM for dipoles of both radial (Figure 3 (a)) and tangential (Figure 4 (a)) directions and all six eccentricities. In the radial direction, the hybrid BE–FE has a maximum RDM of 0.0639±0.0189at 98% source eccentricity, and the PI-FEM has its maximum RDM of 0.0955±0.0405at the same source eccentricity (Figure 3 (a)). Also, the PI-FEM leads to a larger RDM variance than the hybrid BE–FE method. In the tangential direction, the maximum RDM obtained from the hybrid BE–FE method is 0.049±0.012at 98% source eccentricity, while the FEM is higher RDM (0.0902±0.0508) at these eccentricities. On the other hand, the maximum RDM obtained from the PI-FEM is 0.1004±0.0295at 90% source eccentricity (Figure 4 (a)).

For dipoles of radial directions, the hybrid BE–FE method significantly outperforms the BEM. The hybrid BE–FE method has a maximum RDM of 0.0639±0.0189at 98% source eccentricity and the BEM has its maximum RDM of 0.2831±0.0725at the same source eccentricity (Figure 3 (a)). In the tangential direction, the BEM outperforms both the hybrid BE–FE method and PI-FEM at 50%, 60% and 70% source eccentricities. However, its error increased and more than other approaches at 80%, 90% and 98% source eccentricities. The RDM obtained from the hybrid BE–FE method is 0.049±0.012 at 98% source eccentricity, while the BEM has a maximum RDM 0.1388±0.0669 at 80% source eccentricities (Figure 4 (a)).

With regard to the MAG (Figure 3 (b) and Figure 4 (b)), the hybrid BE–FE method outperforms the PI-FEM and BEM. In the radial direction, the hybrid BE–FE has a maximum MAG of 1.0709±0.0298 at 98% source eccentricity. While the PI-FEM and BEM have maximum MAG error of 1.1856±0.2946 and 0.8178±0.0273 at 90% and 98% source eccentricity, respectively (Figure 3 (b)). In the tangential direction, the worst result of MAG from the hybrid BE–FE method is 1.0203±0.0006 at 80% source eccentricity while the PI-FEM is higher MAG (1.1979±0.2284) at these eccentricities. On the other hand, the maximum MAG obtained from the BEM is 1.0423±0.019 at 90% source eccentricity (Figure 4 (b)). Also, the PI-FEM leads to the largest MAG variance with P-value <0.05 at all source eccentricities, as shown in Table 2.

### 4-2 Example II: Anisotropic piece-wise homogenous three-layer spherical head model

For modeling anisotropicity in the EEG forward problem, the hybrid BE–FE method offers an alternative solution. So to assess the performance of the hybrid BE–FE method for the anisotropic three-layer spherical head model, we compared its performance with that of PI-FEM. The radius and conductivity of each layer were the same as those in Table 1, but the conductivity of the skull was anisotropic with an optimized anisotropy ratio 0.0093: 0.015 [30]. It is noteworthy that in this model, the brain (containing dipoles) is modeled by using the BEM, while other layers are modeled by using the FEM.

Figure 5 and Figure 6 show the resulting RDM and MAG for various dipole eccentricities when dipoles are radial and tangential, respectively. Also, the mean and standard deviation of RDM and MAG, and P-values of the Wilcoxon signed-rank test are reported in Table 3. It should be noted that some datasets did not pass the Gaussian test (P-value<0.05). For this reason, we used Wilcoxon signed-rank test to calculate P-value. The significant differences are shown as gray in Table 3. The numbers of nodes, tetrahedral volume elements and triangular surface elements of volume mesh are the same as the previous simulation in section 4-1.

**Figure 5:**
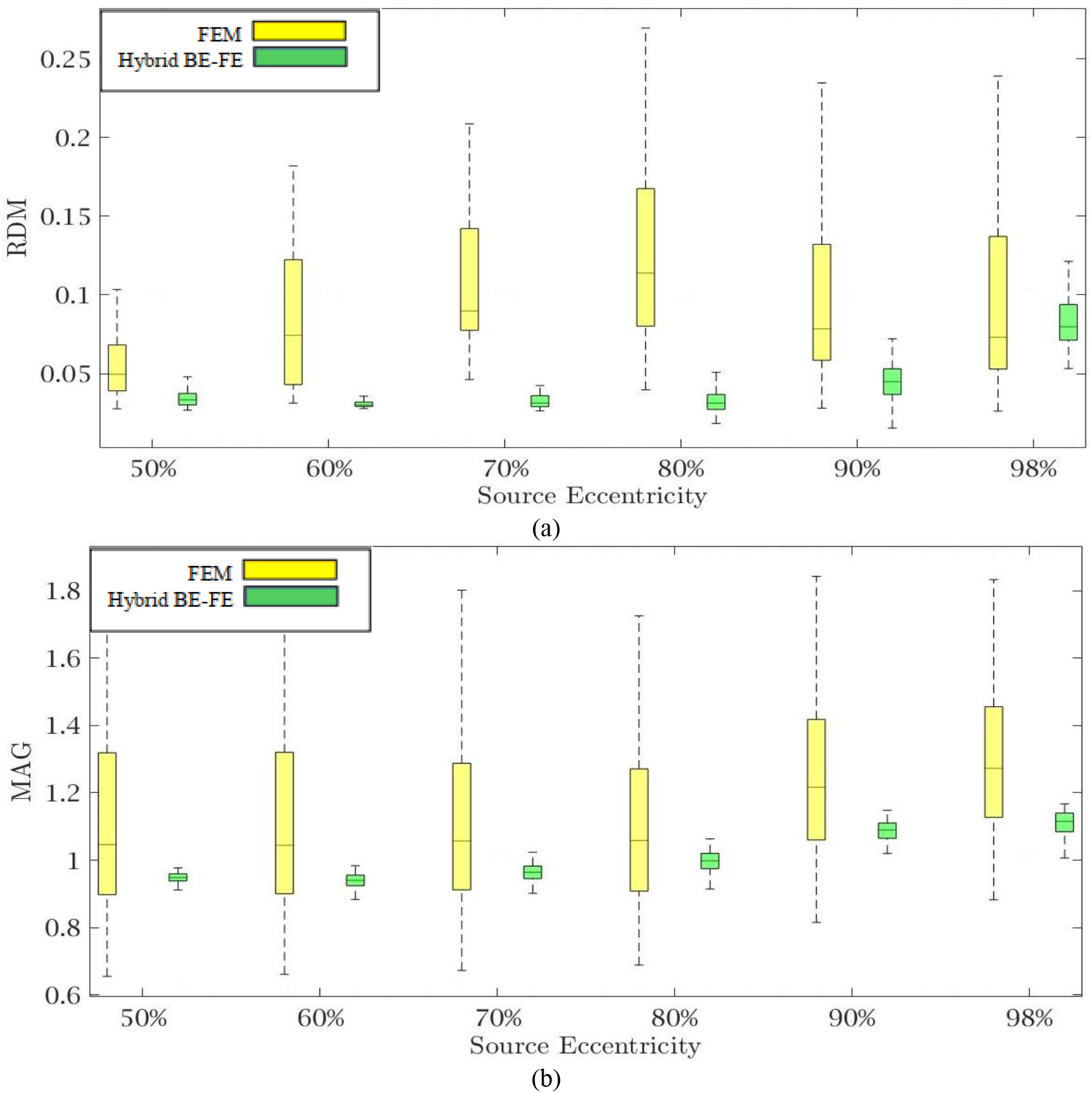
Example II: Anisotropic piece-wise homogenous three-layer spherical head model for radial dipole orientation (z-axis), (a) RDM and (b) MAG boxplots of PI-FEM and hybrid BE–FE methods at six different source eccentricities.

**Figure 6:**
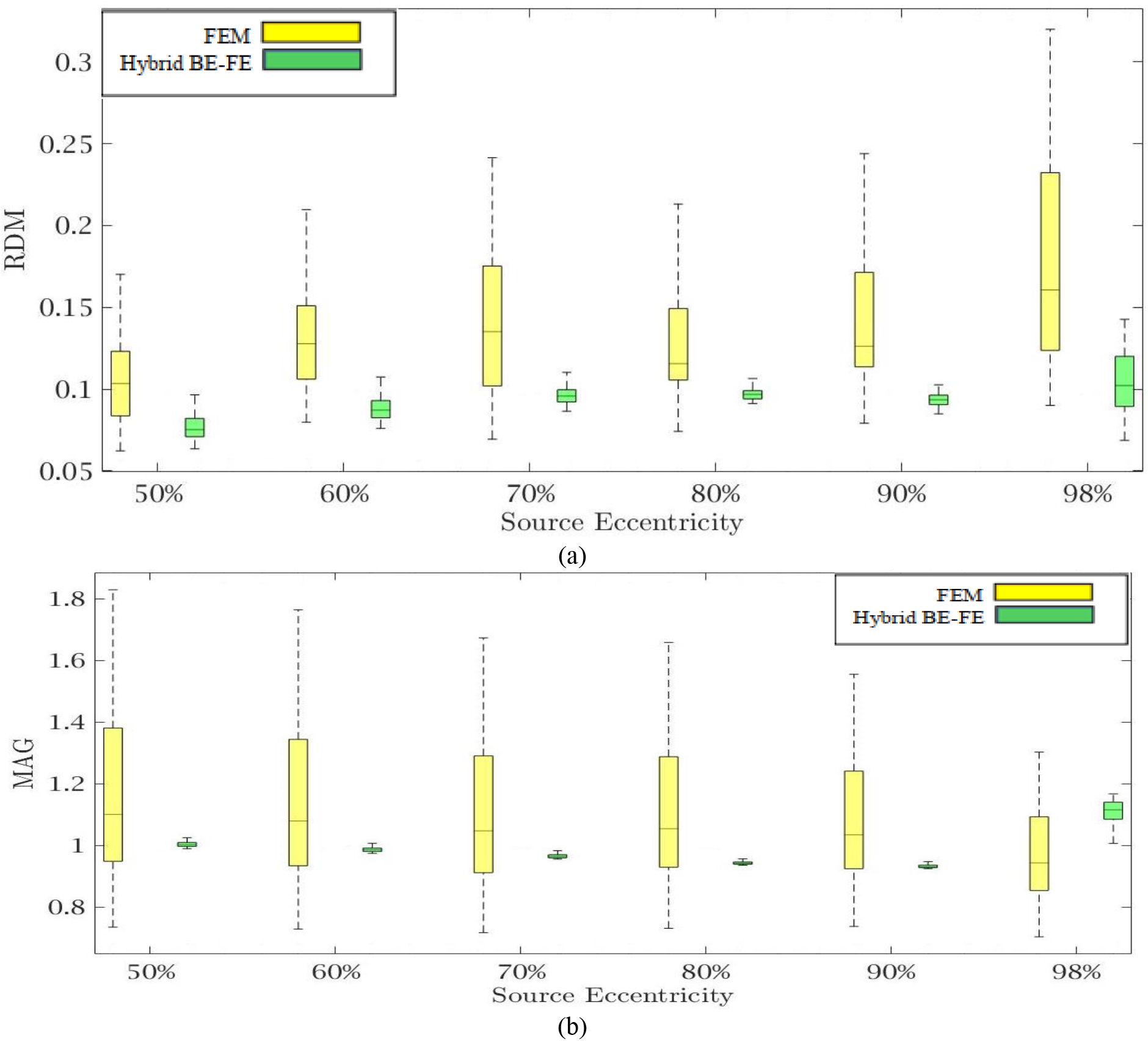
Example II: Anisotropic piece-wise homogenous three-layer spherical head model for tangential dipole orientation (x-axis) (a) RDM and (b) MAG boxplots of PI-FEM and hybrid BE–FE methods at six different source eccentricities.

**Table 3:** Example II: Anisotropic piece-wise homogeneous three-layer spherical head model, mean±std and P-value of Wilcoxon signed-rank test of 50 realizations of RDM and MAG obtained from PI-FEM and hybrid BE–FE method for dipoles of six different source eccentricities and both radial and tangential directions. The significant results are shown as gray.

|  |  | Radial Direction (z-axis) |  |  |  |  |  | Tangential Direction (x-axis) |  |  |  |  |  |
| --- | --- | --- | --- | --- | --- | --- | --- | --- | --- | --- | --- | --- | --- |
| Source Eccentricity |  | 50% | 60% | 70% | 80% | 90% | 98% | 50% | 60% | 70% | 80% | 90% | 98% |
| RDM | PI-FEM | 0.0562<br>$\pm 0.0206$ | 0.0843<br>$\pm 0.0422$ | 0.1059<br>$\pm 0.0395$ | 0.126<br>$\pm 0.0548$ | 0.096<br>$\pm 0.0502$ | 0.0917<br>$\pm 0.0501$ | 0.105<br>$\pm 0.0265$ | 0.1312<br>$\pm 0.0314$ | 0.1411<br>$\pm 0.0426$ | 0.1266<br>$\pm 0.0306$ | 0.1399<br>$\pm 0.0376$ | 0.1762<br>$\pm 0.0594$ |
| | Hybrid BE-FE | 0.0356<br>$\pm 0.0084$ | 0.0323<br>$\pm 0.0061$ | 0.0344<br>$\pm 0.0087$ | 0.0322<br>$\pm 0.0075$ | 0.0437<br>$\pm 0.014$ | 0.08326<br>$\pm 0.0178$ | 0.0785<br>$\pm 0.0126$ | 0.0901<br>$\pm 0.012$ | 0.098<br>$\pm 0.0098$ | 0.0979<br>$\pm 0.006$ | 0.094<br>$\pm 0.0051$ | 0.1158<br>$\pm 0.0474$ |
| | P-value | 1.1913<br>$\times 10^{-7}$ | 8.0311<br>$\times 10^{-10}$ | 9.0681<br>$\times 10^{-10}$ | 7.5569<br>$\times 10^{-10}$ | 1.3025<br>$\times 10^{-9}$ | 0.6887 | 0.237 | 1.0717<br>$\times 10^{-7}$ | 2.0113<br>$\times 10^{-7}$ | 2.742<br>$\times 10^{-7}$ | 7.5824<br>$\times 10^{-9}$ | 1.0872<br>$\times 10^{-9}$ |
| MAG | PI-FEM | 1.1072<br>$\pm 0.273$ | 1.1035<br>$\pm 0.2646$ | 1.105<br>$\pm 0.2517$ | 1.0996<br>$\pm 0.2326$ | 1.2453<br>$\pm 0.2309$ | 1.2934<br>$\pm 0.2143$ | 1.1564±<br>0.2656 | 1.1317<br>$\pm 0.2517$ | 1.0951<br>$\pm 0.233$ | 1.0993<br>$\pm 0.2143$ | 1.0745<br>$\pm 0.188$ | 0.9709<br>$\pm 0.1446$ |
| | Hybrid BE-FE | 0.9474<br>$\pm 0.016$ | 0.9372<br>$\pm 0.0255$ | 0.9632<br>$\pm 0.0289$ | 0.9931<br>$\pm 0.0392$ | 1.0801<br>$\pm 0.0442$ | 1.0925<br>$\pm 0.0722$ | 1.0072±<br>0.0168 | 0.9901<br>$\pm 0.0159$ | 0.9683<br>$\pm 0.0143$ | 0.9454<br>$\pm 0.0115$ | 0.9333<br>$\pm 0.0087$ | 1.0925<br>$\pm 0.0722$ |
| | P-value | 0.0007 | 0.0002 | 0.0014 | 0.0104 | 4.348<br>$\times 10^{-5}$ | 2.4737<br>$\times 10^{-7}$ | 0.0006 | 0.0026 | 0.0169 | 0.0088 | 0.0219 | 0.1333 |

Comparing the hybrid BE–FE method and PI-FEM with regard to the RDM (Figure 5 (a) and Figure 6 (a)) shows the hybrid BE–FE method outperforms the PI-FEM in both directions, especially in the radial direction (P-value<0.05) (Figure 5 (a)). For radial dipoles, the maximum RDM obtained from the hybrid BE–FE method is 0.08326±0.0178at 98% source eccentricity, while the PI-FEM is higher RDM (0.0917±0.0501) at this eccentricity. While the PI-FEM has a maximum RDM error of 0.126±0.0548 at 80% source eccentricity (Figure 5 (a)). For tangential dipoles, the PI-FEM has a maximum RDM of 0.1762±0.0594at 98% source eccentricity while the hybrid BE–FE method leads to a maximum RDM of 0.1158±0.0474at the same source eccentricity. The variance of RDM obtained from the PI-FEM is much greater than that of the hybrid BE–FE method, with P-value <0.05 for all source eccentricities.

The results of MAG (Figure 5 (b) and Figure 6 (b)) clearly show that the hybrid BE–FE method outperforms the PI-FEM for dipoles of both directions. The variance of MAG obtained from the PI-FEM is much greater than that of the hybrid BE–FE method with P-value <0.05 for all source eccentricities.

### 4-3 Example III: Anisotropic piece-wise homogenous four-layer spherical model

We simulate the anisotropic piece-wise homogenous four-layer spherical head model to assess the performance of the hybrid BE–FE method compared with the PI-FEM when a fourth layer (CSF) was considered.

Figure 7 indicates the piece-wise homogeneous four-layer spherical head model. The radius and conductivity interval of these four layers are indicated in Table 4. In this simulation, the PI-FEM mesh has 14350 nodes and 81942 tetrahedral volume elements, and the hybrid BE–FE mesh has 7248 nodes, 33855 tetrahedral volume elements and 3550 triangular surface elements.

**Figure 7:**
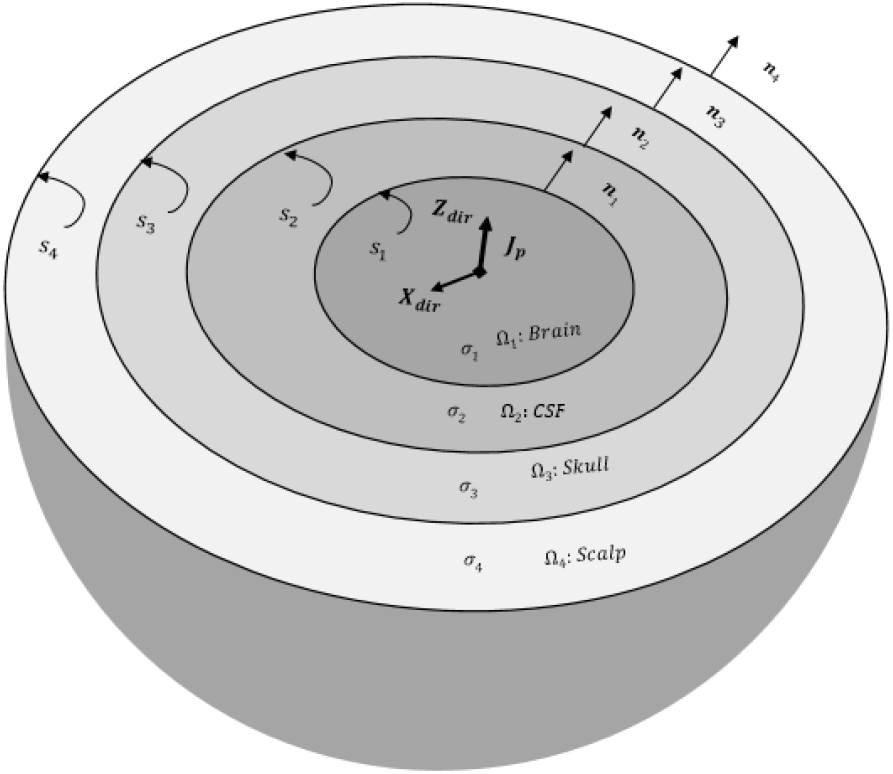
The piece-wise homogenous four-layer spherical head model (brain, CSF, skull, and scalp). The conductivity of the skull can be isotropic or anisotropic.

**Table 4:** Example III: Parameters of the concentric four-layer spherical head model [31], [38]. The conductivity values, radius and optimized anisotropy ratio are based on [31], [38] and [30], respectively.

| Tissue | Brain | CSF | Skull | Scalp |
| --- | --- | --- | --- | --- |
| Outer shell radius (cm) | 7.6 | 8.0 | 8.6 | 9.2 |
| Conductivity interval (S/m) | 0.2200-0.6700 | 1.7696-1.8104 | 0.0016-0.0330 | 0.2800-0.8700 |
| Optimized anisotropy ratio | - | - | 0.0093: 0.015 | - |

Figure 8 and Figure 9 show boxplots of RDM and MAG of the PI-FEM and the hybrid BE–FE method for anisotropic and piece-wise homogeneous four-layer spherical head model (Figure 7) versus different eccentricities of the dipole for radial and tangential dipoles, respectively. For each model, 50 realizations were simulated by randomly chosen conductivities from intervals shown in Table 4. Also, the mean and standard deviation of RDM and MAG and P-values of Wilcoxon signed-rank tests are reported in Table 5. It should be noted that some datasets did not the pass Gaussian test (P-value<0.05). For this reason, we used the Wilcoxon signed-rank test to calculate P-value. The significant results in this table are shown as gray.

**Figure 8:**
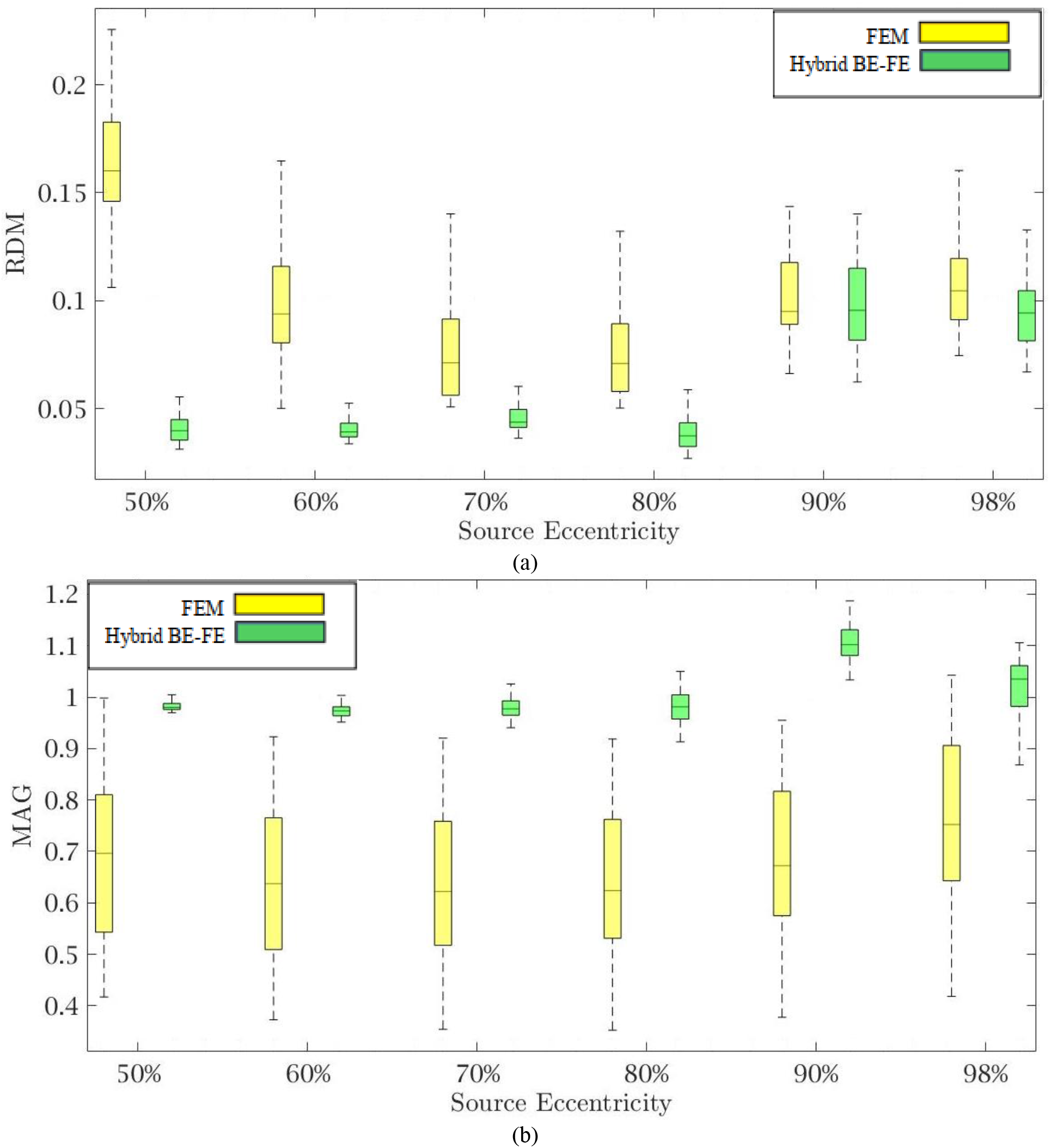
Example III: Anisotropic piece-wise homogenous four-layer spherical head model for radial dipole orientation (z-axis) (a) RDM and (b) MAG boxplots of PI-FEM and hybrid BE–FE methods at six different source eccentricities.

**Table 5:**
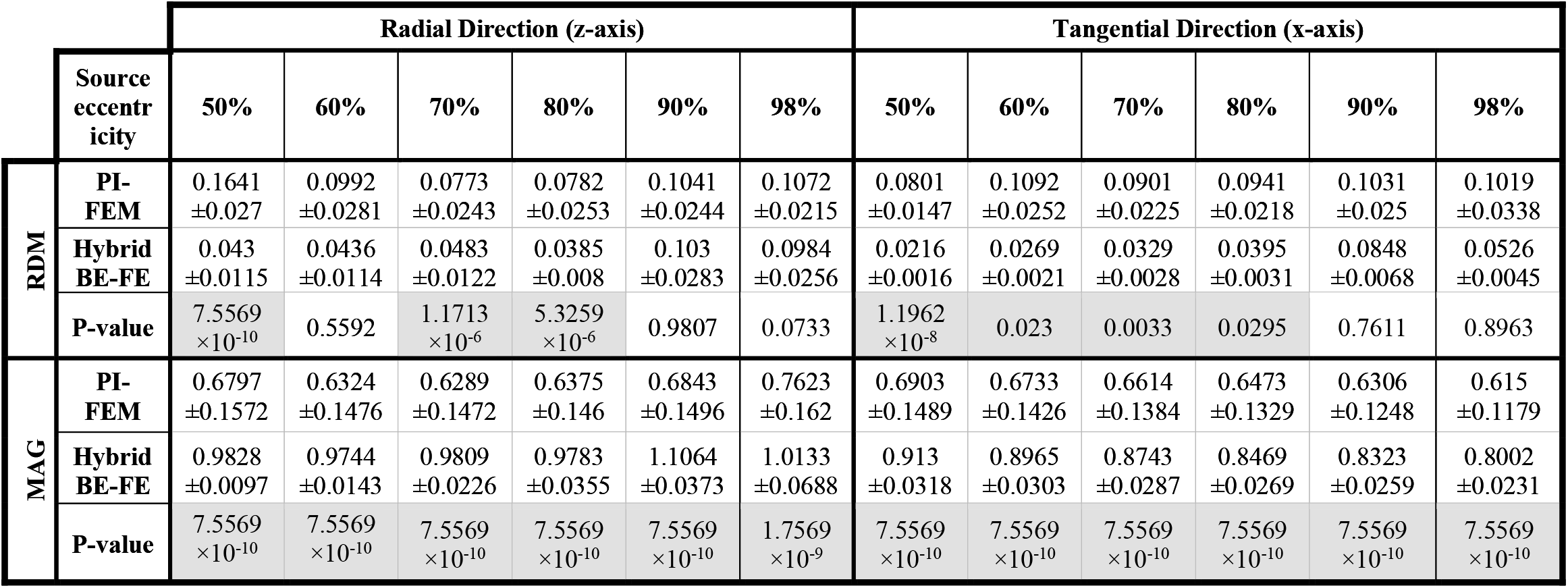
Example III: Anisotropic piece-wise homogeneous four-layer spherical head model, mean±std and P-value of Wilcoxon signed-rank test of 50 realizations of RDM and MAG obtained from PI-FE and hybrid BE–FE methods for dipoles of six different source eccentricities and both radial and tangential directions. The significant results are shown as gray.

|  |  | Radial Direction (z-axis) |  |  |  |  |  | Tangential Direction (x-axis) |  |  |  |  |  |
| --- | --- | --- | --- | --- | --- | --- | --- | --- | --- | --- | --- | --- | --- |
|  | Source eccentricity | 50% | 60% | 70% | 80% | 90% | 98% | 50% | 60% | 70% | 80% | 90% | 98% |
| RDM | PI-FEM | 0.1641<br>$\pm 0.027$ | 0.0992<br>$\pm 0.0281$ | 0.0773<br>$\pm 0.0243$ | 0.0782<br>$\pm 0.0253$ | 0.1041<br>$\pm 0.0244$ | 0.1072<br>$\pm 0.0215$ | 0.0801<br>$\pm 0.0147$ | 0.1092<br>$\pm 0.0252$ | 0.0901<br>$\pm 0.0225$ | 0.0941<br>$\pm 0.0218$ | 0.1031<br>$\pm 0.025$ | 0.1019<br>$\pm 0.0338$ |
| | Hybrid BE-FE | 0.043<br>$\pm 0.0115$ | 0.0436<br>$\pm 0.0114$ | 0.0483<br>$\pm 0.0122$ | 0.0385<br>$\pm 0.008$ | 0.103<br>$\pm 0.0283$ | 0.0984<br>$\pm 0.0256$ | 0.0216<br>$\pm 0.0016$ | 0.0269<br>$\pm 0.0021$ | 0.0329<br>$\pm 0.0028$ | 0.0395<br>$\pm 0.0031$ | 0.0848<br>$\pm 0.0068$ | 0.0526<br>$\pm 0.0045$ |
| | P-value | 7.5569<br>$\times 10^{-10}$ | 0.5592 | 1.1713<br>$\times 10^{-6}$ | 5.3259<br>$\times 10^{-6}$ | 0.9807 | 0.0733 | 1.1962<br>$\times 10^{-8}$ | 0.023 | 0.0033 | 0.0295 | 0.7611 | 0.8963 |
| MAG | PI-FEM | 0.6797<br>$\pm 0.1572$ | 0.6324<br>$\pm 0.1476$ | 0.6289<br>$\pm 0.1472$ | 0.6375<br>$\pm 0.146$ | 0.6843<br>$\pm 0.1496$ | 0.7623<br>$\pm 0.162$ | 0.6903<br>$\pm 0.1489$ | 0.6733<br>$\pm 0.1426$ | 0.6614<br>$\pm 0.1384$ | 0.6473<br>$\pm 0.1329$ | 0.6306<br>$\pm 0.1248$ | 0.615<br>$\pm 0.1179$ |
| | Hybrid BE-FE | 0.9828<br>$\pm 0.0097$ | 0.9744<br>$\pm 0.0143$ | 0.9809<br>$\pm 0.0226$ | 0.9783<br>$\pm 0.0355$ | 1.1064<br>$\pm 0.0373$ | 1.0133<br>$\pm 0.0688$ | 0.913<br>$\pm 0.0318$ | 0.8965<br>$\pm 0.0303$ | 0.8743<br>$\pm 0.0287$ | 0.8469<br>$\pm 0.0269$ | 0.8323<br>$\pm 0.0259$ | 0.8002<br>$\pm 0.0231$ |
| | P-value | 7.5569<br>$\times 10^{-10}$ | 7.5569<br>$\times 10^{-10}$ | 7.5569<br>$\times 10^{-10}$ | 7.5569<br>$\times 10^{-10}$ | 7.5569<br>$\times 10^{-10}$ | 1.7569<br>$\times 10^{-9}$ | 7.5569<br>$\times 10^{-10}$ | 7.5569<br>$\times 10^{-10}$ | 7.5569<br>$\times 10^{-10}$ | 7.5569<br>$\times 10^{-10}$ | 7.5569<br>$\times 10^{-10}$ | 7.5569<br>$\times 10^{-10}$ |

As shown in Figure 8 (a) and Figure 9 (a), with regard to the RDM, the hybrid BE–FE method is more accurate than PI-FEM for both dipole directions. In fact, the mean RDM obtained from the hybrid BE–FE method is significantly smaller (P-value<0.05) than that of PI-FEM for both dipole directions and at most of the eccentricities. The maximum RDM obtained from the PI-FEM is 0.1641±0.027 at 50% source eccentricity at the radial direction, whereas for the hybrid BE–FE method, this value at this source eccentricity is 0.043±0.0115. On the other hand, at the tangential direction (Figure 9 (a)), the mean RDM obtained from the PI-FEM has a maximum of 0.1092±0.0252 at 60% source eccentricity, while the hybrid BE–FE method has a maximum RDM of 0.0848±0.0068 at 90% source eccentricity.

**Figure 9:**
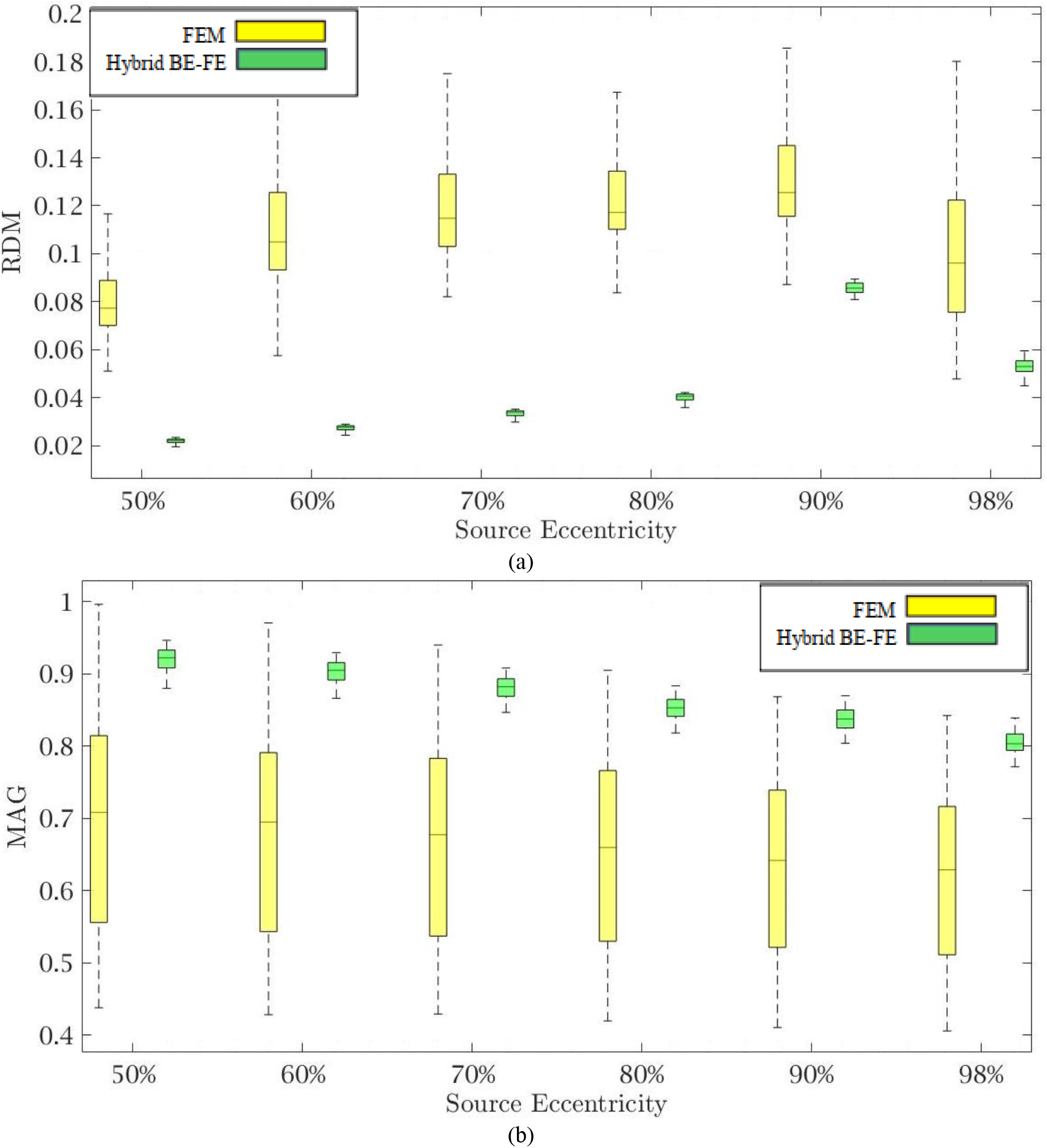
Example III: Anisotropic piece-wise homogenous four-layer spherical head model for tangential dipole orientation (x-axis) (a) RDM and (b) MAG boxplots of PI-FEM and hybrid BE–FE methods at six different source eccentricities.

Also, with regard to the MAG (Figure 8 (b) and Figure 9 (b)), the influence of considering the CSF layer to the spherical head model is apparent. As shown in Figure 8 (b) and Figure 9 (b), the mean MAG obtained from the PI-FEM is highly decreased by considering the fourth layer. On the other hand, the MAG error obtained from the hybrid BE–FE method is significantly much better than PI-FEM (P-value<0.05) in both directions. For radial dipoles, the best results of MAG for the PI-FEM and the hybrid BE–FE method are 0.7623±0.162at 98% source eccentricity and 1.0133±0.0688at the same source eccentricity. On the other hand, for tangential dipoles, the mean MAG obtained from the hybrid BE–FE method is significantly better (P-value<0.05) than that of PI-FEM at all eccentricities. Also, the variance of MAG obtained from the PI-FEM is much bigger than that of the hybrid BE-FE method, which implies the hybrid BE–FE method to be more precise than PI-FEM for tangential dipoles.

### 4-4 Example IV: Anisotropic piece-wise homogenous four-layer realistic head model

Although spherical head models (and associated analytical solutions) are fundamental for reliable assessment of any proposed forward solution strategy, it is of fundamental importance to validate the applicability and performance of the technique proposed here on realistic MRI-obtained head models (Figure 10). These models allow for an individual-based head model to be used in solving the forward problem and result in more precise source localization. The values of conductivity of brain, CSF and scalp are 0.33, 1.79 and 0.43 S/m, respectively [31]. Although the three-layer heterogeneous skull model is more accurate than a single-layer homogenous and anisotropic the skull, we used the simplified model of the skull as a single-layer homogenous and anisotropic profile obtained by automatic segmentation of FieldTrip software [37]. However, our method is able to model a complex profile of the skull when it is available. We use optimized anisotropy ratio 0093/0.015 S/m for skull in this study [30].

**Figure 10:**
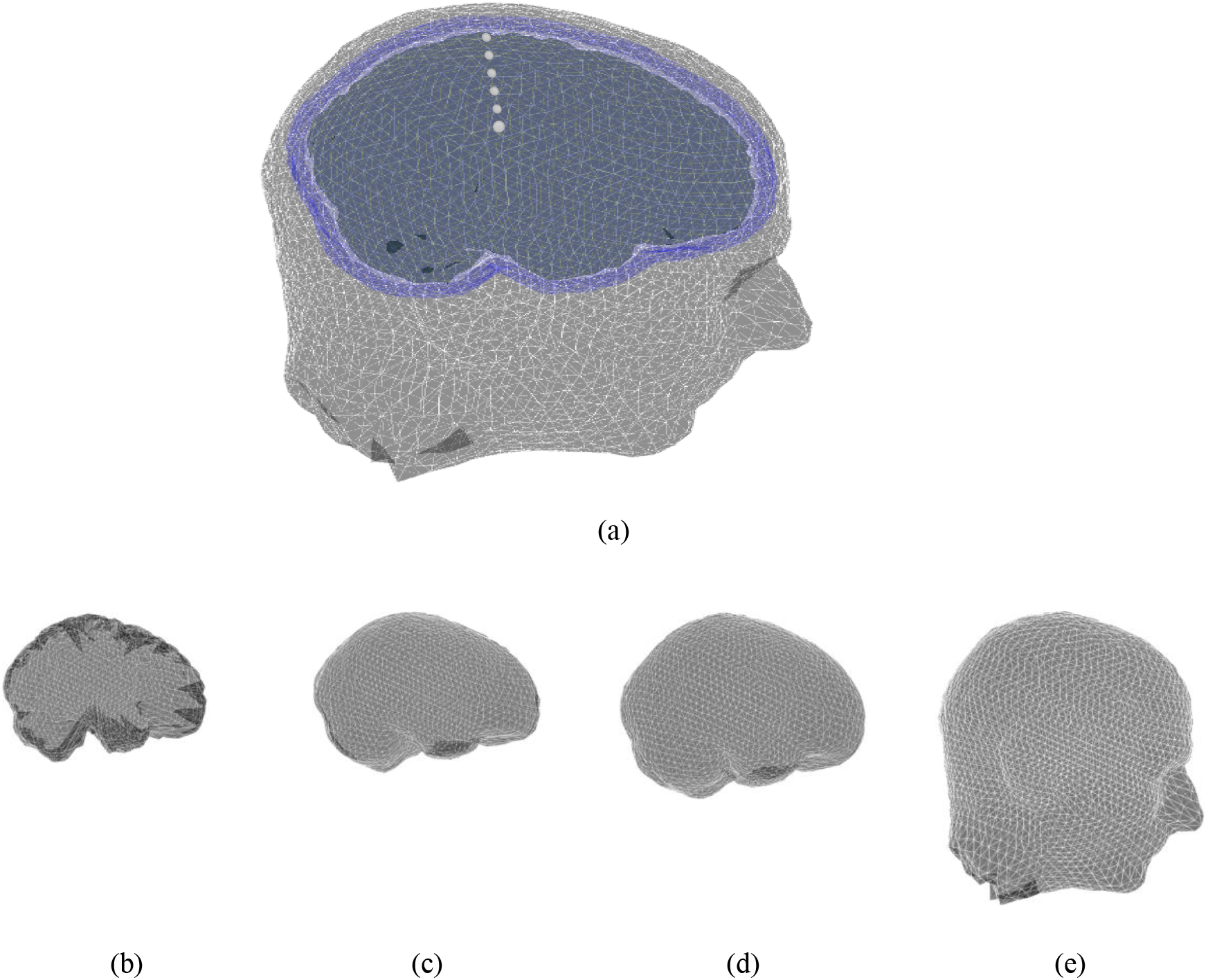
(a) MRI-based realistic head model with four layers: (b) brain, (c) CSF, (d) skull and (e) scalp.

In the realistic head model, the PI-FEM mesh has 221779 nodes and 1380065 tetrahedral volume elements, and the hybrid BE–FE mesh has 108623 nodes, 655756 tetrahedral volume elements and 7004 triangular surface elements.

Since the analytical solution is unavailable in this case, the PI-FEM served as the reference method. The forward problem was solved by using the PI-FEM on a refined model having 10850052 tetrahedrons and 1717389 nodes.

The results of this benchmarking can be seen in Figure 11 and Figure 12 for radial and tangential dipoles, respectively, where a dipolar source was moved from 50% to 98% of the source eccentricity from the center to the surface of the brain, as shown in Figure 10.

**Figure 11:**
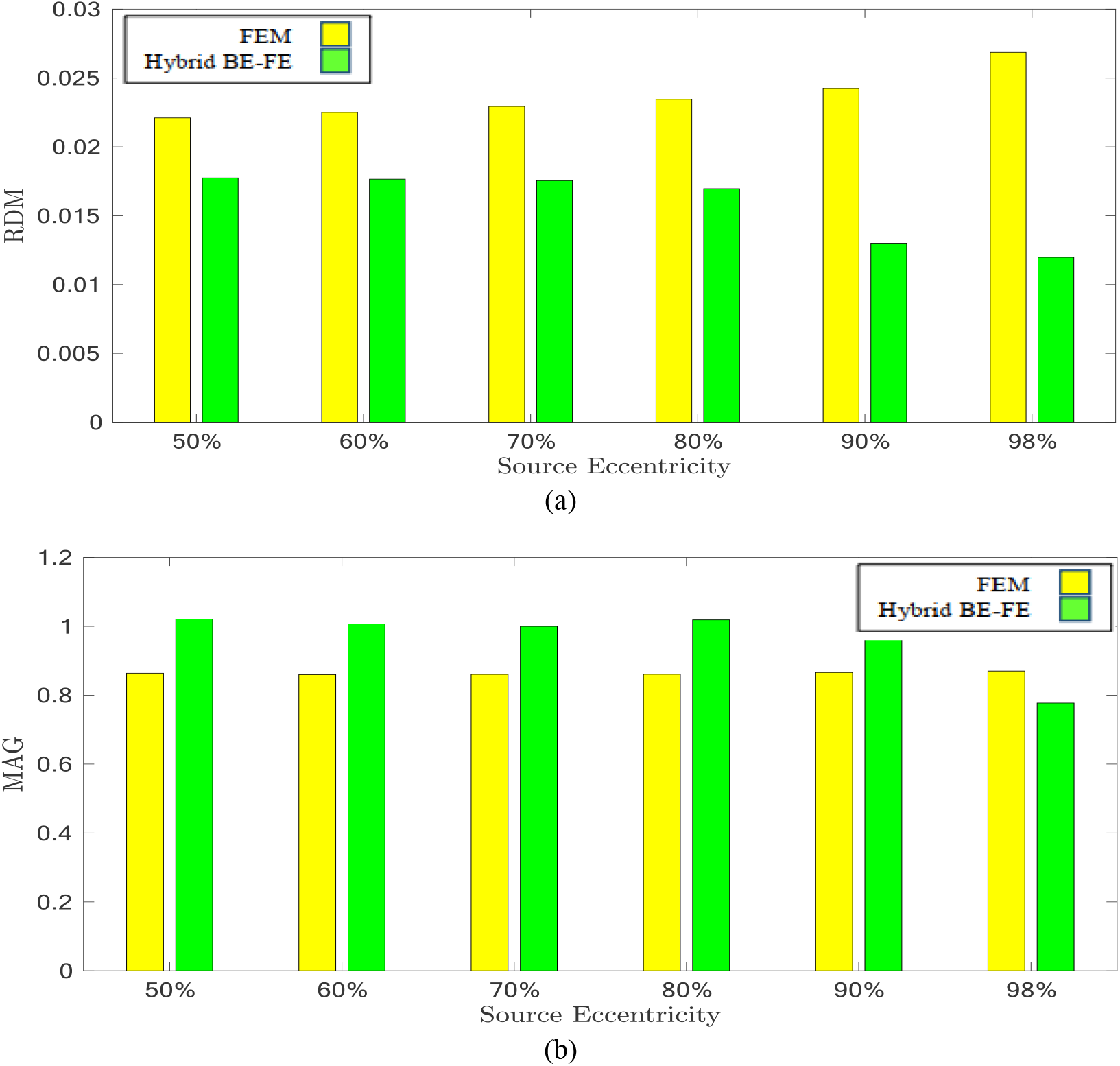
Example IV: Anisotropic piece-wise homogenous four-layer realistic head model for radial dipole orientation (z-axis), (a) RDM and (b) MAG boxplots of PI-FEM and hybrid BE–FE methods at six different source eccentricities.

The RDM and MAG obtained from the hybrid BE–FE method, and PI-FEM presented in this section have been computed with respect to the reference solution.

The hybrid BE–FE method outperforms the PI-FEM with regard to the RDM in both directions (Figure 11 (a) and Figure 12 (a)). The highest RDM values obtained from the hybrid BE–FE method and PI-FEM, for radial dipoles, are 0.0177 at 50% source eccentricity and 0.02686 at 98% source eccentricity, respectively, and for tangential dipoles, are 0.0187 and 0.0245 respectively, both at 98% source eccentricity.

**Figure 12:**
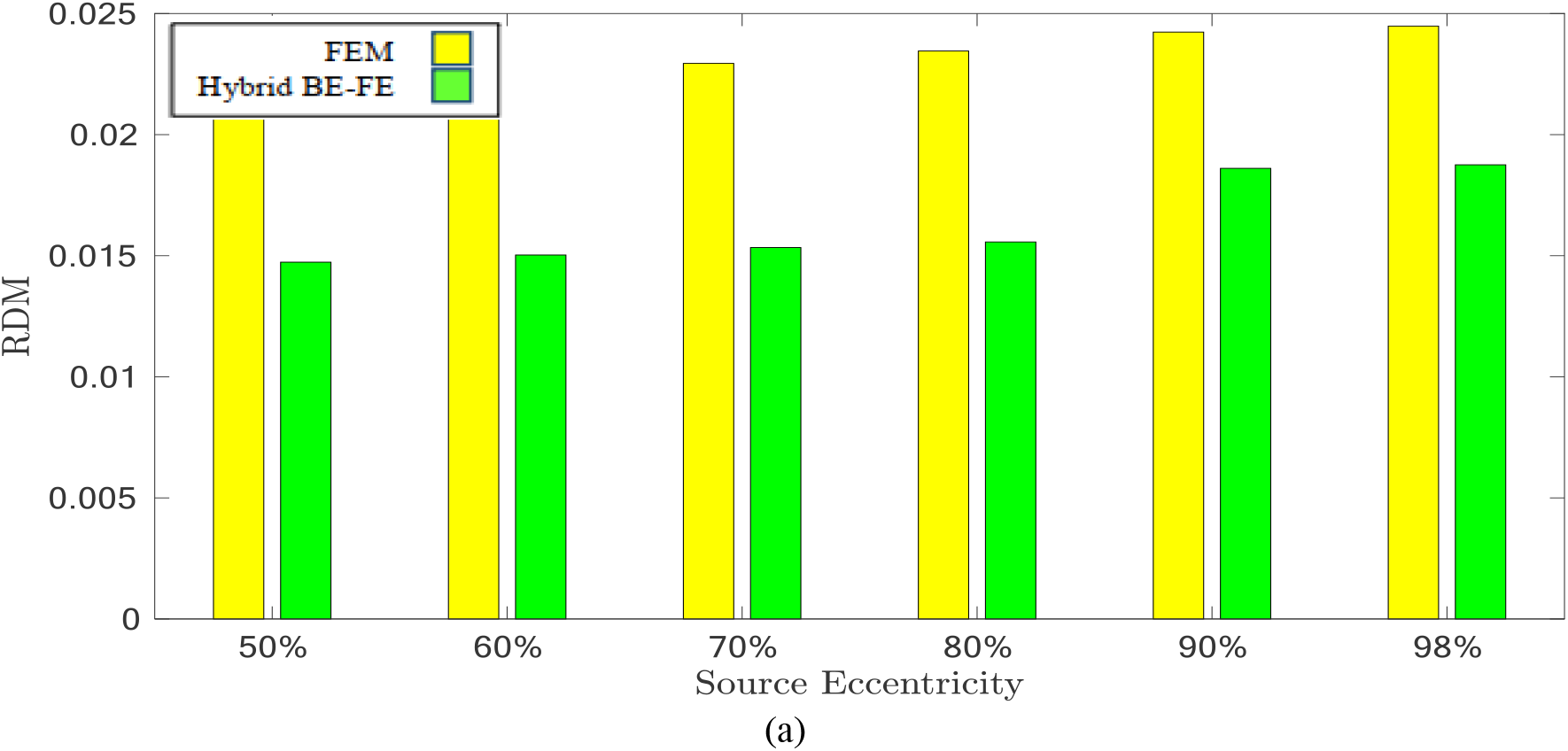
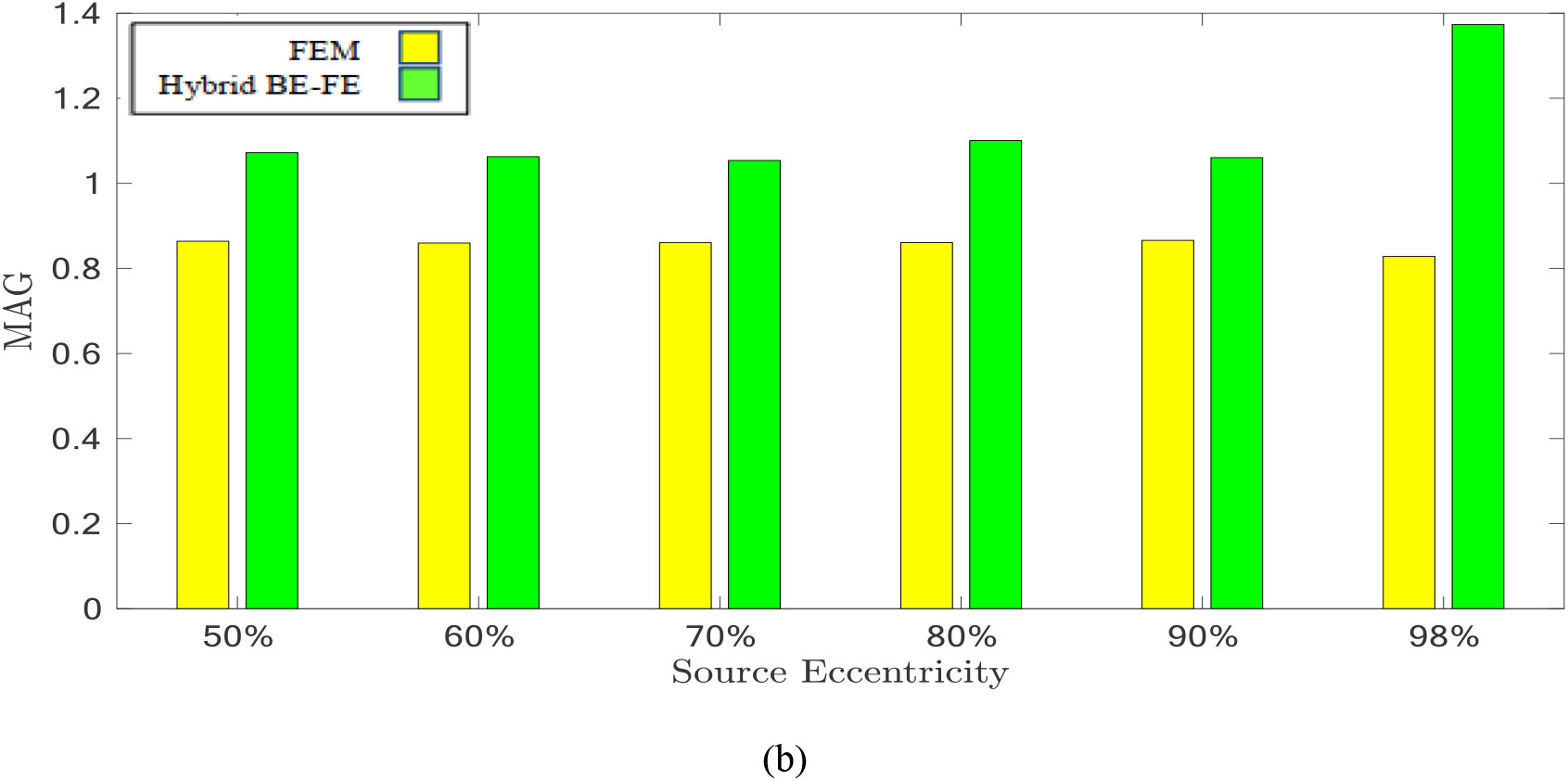
Example IV: Anisotropic piece-wise homogenous four-layer realistic head model for tangential dipole orientation (x-axis) (a) RDM and (b) MAG boxplots of PI-FEM and hybrid BE–FE methods at six different source eccentricities.

The MAG analysis (Figure 11 (b) and Figure 12 (b)) showed the same results as the RDM except in 98%.. The worst MAG values of the hybrid BE–FE method and PI-FEM are, respectively, 0.7771 at 98% source eccentricity and 0.8598 at 60% source eccentricity for radial dipoles, and 1.3734and 0.8284at 98% source eccentricity for tangential dipoles. These results may only be seen as hints since no exact solution exists that can be taken as reference.

## 5 Discussion

Although each BEM and FEM have several advantages, they have drawbacks in solving EEG forward problem. BEM cannot model complex geometry such as inhomogeneity and anisotropicity and surfaces with holes head model [3], [6]. On the other hand, using the FEM causes singularity in right-hand side of EEG forward equation (1) [3], [7]. The coupling method proposed in [25] for iteratively solving the EEG forward problem has two major drawbacks. First, it is very time-consuming to achieve the relative residuals below a properly set value. Second, to solve the global system iteratively, relaxation parameters need to be set at the interfaces to ensure convergence. These relaxation parameters are set manually, and inappropriate values of these parameters would make the scheme diverge. The values of relaxation parameters are not accurate. For using the advantages of both methods, the hybrid BE–FE method offers an alternative solution. The idea behind the hybrid BE–FE method has already been proposed for solving EIT forward problem [26]. In this study, we proposed reformulating the hybrid BE– FE method for solving the EEG forward problem. The hybrid BE–FE method provides an elegant solution to the practical problem of EEG forward problem of how to model heterogeneity and anisotropicity in tissues whose boundaries are known, without complex volume meshing of the whole 3D domain. It is noteworthy that in this method, the volume meshing is not eliminated, but it is limited to regions without dipoles.

Using the PI-FEM, BEM, and hybrid BE–FE method, we compared simulated results for spherical and realistic head models at six different dipole eccentricities and for radial and tangential dipoles. In this study, a layer homogenous isotropic/anisotropic skull model with optimized value was used.

Regarding RDM and MAG, the results of the isotropic inhomogeneous three-layer spherical head model showed that the hybrid BE–FE method outperforms the BEM, especially at the radial direction (see Figure 3 and Figure 4). However, with regard to RDM, the BEM performs better than the hybrid BE–FE method at 50%, 60% and 70% of source eccentricities in the tangential direction, but it performs the worst at 80%, 90% and 98% of source eccentricities and with regard to MAG the hybrid BE–FE method outperforms the BEM. To be noticed that the BEM cannot simulate inhomogeneous and anisotropic media, the hybrid BE–FE method can be a good alternative to be used instead of the BEM in the multi-layer medium.

In the isotropic three-layer spherical head model, the PI-FEM has the worst performance at 50%, 60% and 70% of source eccentricities in both directions (see Figure 3 and Figure 4). Also, it has worse performance than the hybrid BE–FE method in all eccentricities. Nevertheless, the hybrid BE–FE method clearly performs well. On the other hand, the MAG of the PI-FEM has a large variance in both directions. While the hybrid BE–FE method has the best results in both directions.

By considering the skull as a layer homogenous anisotropic model, results showed that the hybrid BE–FE method outperforms the PI-FEM at all eccentricities in both directions (Figure 5 and Figure 6). In the radial direction, the RDM of the hybrid BE–FE method is much smaller than the PI-FEM. On the other hand, although the difference between their RDM in the tangential direction is not as much as in the radial direction, the RDM of the hybrid BE–FE method is still better than the PI-FEM (see Figure 5 (a) and Figure 6 (a)). Therefore, the hybrid BE–FE method clearly performs well and has small RDM less than 0.08326 for radial direction and 0.1158 for tangential direction. While the PI-FEM has a maximum RDM of 0.126 and 0.1762 in radial and tangential direction, respectively (Table 3). On the other hand, the MAG of the PI-FEM has a large variance in both directions (see Figure 5 (b) and Figure 6 (b)). While the hybrid BE–FE method has the best results in both directions.

In the next step, we compared the PI-FEM and hybrid BE–FE method when a fourth layer (CSF) was considered. With regard to the RDM, in radial direction, the hybrid BE–FE method is more accurate than the PI-FEM (see Table 5). With regard to the MAG, the hybrid BE–FE method performs clearly better than the PI-FEM. At the tangential direction, the hybrid BE–FE method outperforms the PI-FEM with regard to the RDM and MAG.

The results of the spherical head model show that the hybrid BE–FE method has higher accuracy than the PI-FEM in both directions. Also, the variance of the PI-FEM is very higher than the hybrid BE–FE method. It shows that by variation of conductivity, the performance of the hybrid BE–FE method is more stable than the PI-FEM. The comparison with the hybrid BE–FE method and PI-FEM for dipoles of different orientations and eccentricities showed that the hybrid BE–FE method leads to higher accuracy.

The result of the realistic head model shows that the hybrid BE–FE method outperforms the PI-FEM in both directions regarding RDM. Also, with regard to MAG, the hybrid BE–FE method outperforms the PI-FEM in both directions except at 98% source eccentricity. However, no exact solution exists as a reference to conclude about the realistic head model.

The overall higher accuracy of the hybrid BE–FE method was expected due to theoretical considerations behind the hybrid BE–FE method since it uses each of the BEM and FEM on the domains better suited for them. It uses the BEM to model the brain layer containing dipoles to avoid the singularity problem of the FEM. On the other side, it uses the FEM to model inhomogeneous and anisotropic compartments to overcome the BEM disability in modeling inhomogeneity and anisotropicity.

There are some limitations in our study that should be addressed. First, the hybrid BE–FE method is more time consuming than the PI-FEM. For example, in the four-layer spherical head model with the same DOF, in our computer simulation study at hand, on the Microsoft Windows 10 Enterprise N, PC with Intel core i7-4510U 2.6GHz CPU and 6-GB RAM, the total forward simulation time of the hybrid BE–FE method was three times more than the FEM. Also, the extracting mesh algorithm of the hybrid BE–FE method is more complex than FEM. There are some academic software tools that generating volume mesh for the FEM very fast and accurately. But to generate mesh for the hybrid BE–FE method, first, we need to generate volume mesh, then to extract mesh to use in the hybrid BE–FE method. Hence, it is more time consuming and complex than the FEM.

## 6 Conclusion

We presented the theory, verification, and evaluation of a hybrid BE–FE method for solving the EEG forward problem. The simulation results of spherical head models demonstrated that the hybrid BE–FE method is more accurate and precise than the PI-FEM. The error (measured by RDM and MAG) obtained from the hybrid BE–FE method indicated that it performs well for deep dipoles. However, when dipoles became close to the first conductivity jump, the error of the hybrid BE–FE method increases, but still less than the PI-FEM at the same eccentricities.

The EEG forward simulation in the realistic head model showed that the hybrid BE–FE method outperforms the PI-FEM in both directions. Overall, our simulations confirm that the hybrid BE–FE method is a promising new approach for solving heterogeneous isotropic/anisotropic EEG forward problems that can outperform the FEM.

The directions of the future work can include further development of the proposed method in other properties of head layers, such as the radius of layers and different meshes. Also, we have an idea to develop a mesh generation algorithm to be used in well-known academic software packages and to be simulated faster. Moreover, we can compare the FEM and our proposed hybrid BE–FE method for a three-layer heterogeneous skull model and consider the anisotropic profile of white matter. Additionally, developing the methodology for use in some applications, such as increasing the accuracy of source localization to detect some diseases, is an attractive field of future research.

## References

[1] Z. Akalin Acar and S. Makeig, “Effects of forward model errors on EEG source localization,” Brain Topogr., vol. 26, no. 3, pp. 378–396, 2013, doi: 10.1007/s10548-012-0274-6.

[2] D. Güllmar, J. Haueisen, and J. R. Reichenbach, “NeuroImage In fl uence of anisotropic electrical conductivity in white matter tissue on the EEG / MEG forward and inverse solution. A high-resolution whole head simulation study,” Neuroimage, vol. 51, no. 1, pp. 145–163, 2010, doi: 10.1016/j.neuroimage.2010.02.014.

[3] J. C. de Munck, C. H. Wolters, and M. Clerc, “EEG and MEG: forward modeling,” in Handbook of Neural Activity Measurement, no. November 2017, 2012, pp. 192–256.

[4] J. Vorwerk, C. Engwer, S. Pursiainen, and C. H. Wolters, “A Mixed Finite Element Method to Solve the EEG Forward Problem,” IEEE Trans. Med. Imaging, vol. 36, no. 4, pp. 930–941, 2017, doi: 10.1109/TMI.2016.2624634.

[5] M. Stenroos and J. Sarvas, “Bioelectromagnetic forward problem: Isolated source approach revis(it)ed,” Phys. Med. Biol., vol. 57, no. 11, pp. 3517–3535, 2012, doi: 10.1088/0031-9155/57/11/3517.

[6] L. Rahmouni, R. Mitharwal, and F. P. Andriulli, “Two volume integral equations for the inhomogeneous and anisotropic forward problem in electroencephalography,” J. Comput. Phys., vol. 348, pp. 732–743, 2017, doi: 10.1016/j.jcp.2017.07.013.

[7] Y. Zhang, Z. Ren, and D. Lautru, “Finite element modeling of current dipoles using direct and subtraction methods for EEG forward problem,” COMPEL-Int. J. Comput. Math. Electr. Electron. Eng., vol. 33, no. 1-2, pp. 210–223, 2014, doi: 10.1108/COMPEL-11-2012-0329.

[8] L. Beltrachini, “The analytical subtraction approach for solving the forward problem in EEG,” Journal of neural engineering. 2019, Accessed: May 11, 2021. [Online]. Available: https://iopscience.iop.org/article/10.1088/1741-2552/ab2694/meta.

[9] C. H. Wolters, A. Anwander, X. Tricoche, D. Weinstein, M. A. Koch, and R. S. MacLeod, “Influence of tissue conductivity anisotropy on EEG/MEG field and return current computation in a realistic head model: A simulation and visualization study using high-resolution finite element modeling,” Neuroimage, vol. 30, no. 3, pp. 813–826, 2006, doi: 10.1016/j.neuroimage.10.014.

[10] K. A. Awada, D. R. Jackson, J. T. Williams, D. R. Wilton, S. B. Baumann, and A. C. Papanicolaou, “Computational aspects of finite element modeling in EEG source localization,” IEEE Trans. Biomed. Eng., vol. 44, no. 8, pp. 736–752, 1997, doi: 10.1109/10.605431.

[11] M. Darbas and S. Lohrengel, “Review on Mathematical Modelling of Electroencephalography (EEG),” Jahresbericht der Dtsch. Math., vol. 121, no. 1, pp. 3–39, 2019, doi: 10.1365/s13291-018-0183-z.

[12] H. Buchner et al., “Inverse localization of electric dipole current sources in finite element models of the human head,” Electroencephalogr. Clin. Neurophysiol., vol. 102, no. 4, pp. 267–278, 1997, doi: 10.1016/S0013-4694(96)95698-9.

[13] Y. Yan, P. L. Nunez, and R. T. Hart, “Finite-element model of the human head: scalp potentials due to dipole sources,” Med. Biol. Eng. Comput., vol. 29, no. 5, pp. 475–481, 1991, doi: 10.1007/BF02442317.

[14] H. Hallez et al., “Review on solving the forward problem in EEG source analysis,” Journal of NeuroEngineering and Rehabilitation, vol. 4. 2007, doi: 10.1186/1743-0003-4-46.

[15] L. Beltrachini, “A Finite Element Solution of the Forward Problem in EEG for Multipolar Sources,” IEEE Trans. Neural Syst. Rehabil. Eng., vol. 27, no. 3, pp. 368–377, 2019, doi: 10.1109/TNSRE.2018.2886638.

[16] C. H. Wolters, H. Köstler, C. Möller, J. Härdtlein, and A. Anwander, “Numerical approaches for dipole modeling in finite element method based source analysis,” Int. Congr. Ser., vol. 1300, pp. 189–192, 2007, doi: 10.1016/j.ics.2007.02.014.

[17] S. Lew, C. H. Wolters, T. Dierkes, C. Röer, and R. S. MacLeod, “Accuracy and run-time comparison for different potential approaches and iterative solvers in finite element method based EEG source analysis,” Appl. Numer. Math., vol. 59, no. 8, pp. 1970–1988, 2009, doi: 10.1016/j.apnum.2009.02.006.

[18] J. Vorwerk, A. Hanrath, C. H. Wolters, and L. Grasedyck, “The multipole approach for EEG forward modeling using the finite element method,” Neuroimage, vol. 201, pp. 1–28, 2019, doi: 10.1016/j.neuroimage.2019.116039.

[19] T. Medani, D. Lautru, D. Schwartz, Z. Ren, and G. Sou, “FEM method for the EEG forward problem and improvement based on modification of the Saint Venant’s method,” Prog. Electromagn. Res., vol. 153, no. Icm, pp. 11–22, 2015, doi: 10.2528/PIER15050102.

[20] M. Y. Monin, L. Rahmouni, A. Merlini, and F. P. Andriulli, “A Hybrid Volume-Surface-Wire Integral Equation for the Anisotropic Forward Problem in Electroencephalography,” IEEE J. Electromagn. RF Microwaves Med. Biol., vol. 4, no. 4, pp. 286–293, 2020, doi: 10.1109/JERM.2020.2966121.

[21] G. Adde, M. Clerc, O. Faugeras, R. Keriven, J. Kybic, and T. Papadopoulo, “Symmetric BEM formulation for the M/EEG forward problem,” Lect. Notes Comput. Sci. (including Subser. Lect. Notes Artif. Intell. Lect. Notes Bioinformatics), vol. 2732, no. 1, pp. 524–535, 2003, doi: 10.1007/978-3-540-45087-0_44.

[22] C. P. Bradley, G. M. Harris, and A. J. Pullan, “The computational performance of a high-order coupled FEM/BEM procedure in electropotential problems,” IEEE Trans. Biomed. Eng., vol. 48, no. 11, pp. 1238–1250, 2001, doi: 10.1109/10.959319.

[23] J. Sikora, R. Bayford, and L. Horesh, “The application of hybrid BEM/FEM methods to solve electrical impedance tomography’s forward problem for the human head,” XII ICEBI V EIT Conf. Gdansk, 2004.

[24] S. Srinivasan, H. R. Ghadyani, B. W. Pogue, and K. D. Paulsen, “A coupled finite element-boundary element method for modeling Diffusion equation in 3D multi-modality optical imaging,” Biomed. Opt. Express, vol. 1, no. 2, p. 398, 2010, doi: 10.1364/boe.1.000398.

[25] E. Olivi, M. Clerc, and T. Papadopoulo, “Domain decomposition for coupling finite and boundary element methods in EEG,” in IFMBE Proceedings, 2010, vol. 28, pp. 120–123, doi: 10.1007/978-3-642-12197-5_24.

[26] P. Ghaderi Daneshmand and R. Jafari, “A 3D hybrid BE–FE solution to the forward problem of electrical impedance tomography,” Eng. Anal. Bound. Elem., vol. 37, no. 4, pp. 757–764, Apr. 2013, doi: 10.1016/j.enganabound.2013.01.016.

[27] C. Ramon, P. H. Schimpf, and J. Haueisen, “Influence of head models on EEG simulations and inverse source localizations,” Biomed. Eng. Online, vol. 5, pp. 1–13, 2006, doi: 10.1186/1475-925X-5-10.

[28] J. Vorwerk, J. H. Cho, S. Rampp, H. Hamer, T. R. Knösche, and C. H. Wolters, “A guideline for head volume conductor modeling in EEG and MEG,” Neuroimage, vol. 100, pp. 590–607, 2014, doi: 10.1016/j.neuroimage.2014.06.040.

[29] M. Akhtari et al., “Conductivities of three-layer line human skull,” Brain Topogr., vol. 14, no. 3, pp. 151–167, 2002, doi: 10.1023/A:1014590923185.

[30] M. Dannhauer, B. Lanfer, C. H. Wolters, and T. R. Knösche, “Modeling of the human skull in EEG source analysis,” Hum. Brain Mapp., vol. 32, no. 9, pp. 1383–1399, 2011, doi: 10.1002/hbm.21114.

[31] J. Vorwerk, Ü. Aydin, C. H. Wolters, and C. R. Butson, “Influence of head tissue conductivity uncertainties on EEG dipole reconstruction,” Front. Neurosci., vol. 13, no. JUN, pp. 1–17, 2019, doi: 10.3389/fnins.2019.00531.

[32] M. Hämäläinen, R. Hari, R. J. Ilmoniemi, J. Knuutila, and O. V. Lounasmaa, “Magnetoencephalography theory, instrumentation, and applications to noninvasive studies of the working human brain,” Rev. Mod. Phys., vol. 65, no. 2, pp. 413–497, 1993, doi: 10.1103/RevModPhys.65.413.

[33] W. Ang, A Beginner’s Course in Boundary Element Methods. Universal, 2007.

[34] J. Jin, The Finite Element Method in Electromagnetics, Third edit. John Wiley & Sons, 2014.

[35] P. H. Schimpf, C. Ramon, and J. Haueisen, “Dipole models for the EEG and MEG,” IEEE Trans. Biomed. Eng., vol. 49, no. 5, pp. 409–418, 2002, doi: 10.1109/10.995679.

[36] Q. Fang and D. A. Boas, “Tetrahedral mesh generation from volumetric binary and grayscale images,” in Proceedings – 2009 IEEE International Symposium on Biomedical Imaging: From Nano to Macro, ISBI 2009, 2009, pp. 1142–1145, doi: 10.1109/ISBI.2009.5193259.

[37] R. Oostenveld, P. Fries, E. Maris, and J. M. Schoffelen, “FieldTrip: Open source software for advanced analysis of MEG, EEG, and invasive electrophysiological data,” Comput. Intell. Neurosci., vol. 2011, 2011, doi: 10.1155/2011/156869.

[38] S. Wagner et al., “Using reciprocity for relating the simulation of transcranial current stimulation to the EEG forward problem,” Neuroimage, vol. 140, pp. 163–173, 2016, doi: 10.1016/j.neuroimage.2016.04.005.

[39] Z. Zhi, “A fast method to compute surface potentials generated by dipoles within multilayer anisotropic spheres,” Phys. Med. Biol., vol. 40, no. 3, pp. 335–349, 1995, doi: 10.1088/0031-9155/40/3/001.

[40] J. W. H. Mejis, O. W. Weier, M. J. Peters, and A. Van Oosterom, “On the Numerical Accuracy of the Boundary Element Method,” IEEE Trans. Biomed. Eng., vol. 36, no. 10, pp. 1038–1049, 1989, doi: 10.1109/10.40805.

[41] J. Malmivuo and R. Plonsey, Bioelectromagnetism: Principles and Applications of Bioelectric and Biomagnetic Fields. Oxford University Press, USA, 2002.

